# Maize genetic diversity is largely unstructured by human ethnolinguistic diversity in its center of origin

**DOI:** 10.64898/2026.02.02.703385

**Authors:** Samantha Snodgrass, Forrest Li, Sowmya Mambakkam, Santiago G. Medina-Muñoz, Luke Sparreo, Mitra Menon, Claudia Perez, Namrata Kalsi, Amit Gourav Ghosh, Stephan C. Schuster, Hie Lim Kim, Andrés Moreno-Estrada, Daniel Runcie, Graham Coop, Jeffrey Ross-Ibarra

## Abstract

Population structure and environmental features often capture major axes of genetic variation in many species. Yet the impacts of human activity often remain unquantified. For domesticated species that rely on human activity for survival and dispersal, human movements and cultural differences may play key roles patterning genetic diversity. Maize is a staple crop of enormous cultural importance to indigenous peoples of the Americas, but cannot survive or disperse without farmers. Using publicly available genotyping and passport data from approximately 2,000 traditional maize varieties, more than 500 whole genome sequences of humans from Mexico, and indigenous linguistic maps, we quantify anthropogenic effects on maize genetic diversity in the Americas. Maize shows very little overall structure, highlighting the effectiveness of indigenous farmers in moving and mixing maize populations. While principal components of maize diversity show meaningful correlations to human genetic diversity, our linear modeling suggests little additional impact of human population structure beyond shared geography. Though differences in maize diversity are often patterned by language locally, we find only weak genome-wide effects at larger spatial scales. Despite the relatively weak global signal of anthropogenic effects, linguistic GWAS, outlier *F_ST_*analyses, and selection scans identified loci associated with specific languages. Leveraging landscape-level sequencing data, we highlight how anthropogenic factors have shaped patterns of maize genetic diversity across Mesoamerica.

## Introduction

Genetic population structure can arise through isolation by distance, adaptation to different environments, or the interplay of these two processes. These processes can result in correlations between population genetic structure and geography (Blanquart *et al*. 2012; Wang 2013) or other barriers to gene flow (Schweizer *et al*. 2016; Ringbauer *et al*. 2018; Li *et al*. 2019). Domesticated species’ genetic diversity is structured by these processes but also shaped by their history of dynamic cultural and biological interactions with humans, a form of bio-cultural coevolution (Weiss *et al*. 2006; Ullah *et al*. 2015; Zeder 2015; Hendry *et al*. 2017; Angourakis *et al*. 2022; Purugganan 2022; Yoder 2025). These processes continue into the present day, as contemporary communities actively maintain, select, and disperse plant materials in ways that impact crop genetic diversity (Louette *et al*. 1997; Louette and Smale 2000; van Heerwaarden *et al*. 2010; Rojas-Barrera *et al*. 2019).

As a result of this close coevolutionary relationship, both human dispersal and cultural differences are expected to spatially pattern genetic diversity in domesticated species. As farmers disperse, they often bring their domesticates with them (Spengler 2020; Zeder 2008), and thus spatial patterns of domesticate genetic diversity may reflect human population genetic structure. For instance, studies of domesticated species such as ancient dog (Bergström *et al*. 2020a; Zhang *et al*. 2025) and sheep (Daly *et al*. 2025) have demonstrated a mirrored history of migration and admixture to demographic histories of ancient human genomes. Barriers and facilitators of dispersal may not only be physical in nature but can also be cultural, influenced by interactions due to ethnolinguistic boundaries. Recent studies have shown correlations between language contact rates and population structure among human populations (Graff *et al*. 2025). If cultural and linguistic differences impact genetic exchange among human populations, they may serve as barriers to trade and adoption of non-local seed as well (Staller *et al*. 2006). Studies have revealed relationships between human ethnolinguistic patterns and genetic and phenotypic diversity in several crops (e.g. flowering time and pathogen networks in sorghum, pearl and fonio millet Labeyrie *et al*. 2014; Delêtre *et al*. 2021; Naino Jika *et al*. 2017; Abrouk *et al*. 2020; Diop *et al*. 2025) and that individual genetic variants can be associated with linguistic groupings (Abrouk *et al*. 2020).

Domesticated maize (*Zea mays* ssp. *mays*) is an annual crop species with significant cultural and economic importance that was domesticated in modern-day Mexico approximately 10 thousand years before present (Matsuoka *et al*. 2002; Piperno *et al*. 2009). Maize requires human intervention for both seed dispersal and survival, and thus patterns in maize genetic diversity should strongly reflect thousands of years of human interaction. Archaeological studies have supported co-location of human economic specialization with improvement of early maize (Smith 2001), and human activity facilitated the spread of maize far from the tropical lowlands of its wild progenitor *Zea mays* ssp. *parviglumis* (Doebley 2004; Doebley *et al*. 2006). Dispersal into the highlands of Mexico led to adaptive differentiation along elevation, aided in part by repeated adaptive introgression from a second wild relative *Zea mays* ssp. *mexicana* (Janzen *et al*. 2022; Li *et al*. 2025; Mercer *et al*. 2008; Calfee *et al*. 2021; Yang *et al*. 2023). Contemporary maize domestication is ongoing through the selection of seed by modern farmers (Rojas-Barrera *et al*. 2019; Bellon *et al*. 2018), and migration of maize seed is further influenced by farmer decision-making at the market level (Dyer and Taylor 2008), farmer kinship networks (Lugo-Castilla *et al*. 2023), and informal seed exchanges (Fenzi *et al*. 2021), thus potentially generating patterns of genetic diversity that are reflective of the human processes governing maize gene flow (van Heerwaarden *et al*. 2010).

Furthermore, the cultural importance of maize in the Americas may lead to varying patterns of use and trade that differ among indigenous groups. Differences in culinary preference or agricultural methods between ethnolinguistic groups, for example, could affect the maize varieties each group grows (Gilabert *et al*. 2025; Vilela *et al*. 2020). Documentation of these cultural factors, however, is highly heterogeneous, making it difficult to perform large-scale comparative work. Language, however, can serve as a useful proxy for maize-related cultural differences, including group-specific words to describe plant species (Maffi 2005; Luna-José and Aguilar 2012), traditional food preparation and storage methods (Cuj *et al*. 2020; Moreno *et al*. 2006), and local agricultural and soil systems (Leclerc and Coppens d’Eeckenbrugge 2012; Merrill 2012; Bautista and Zinck 2010; León Villalobos *et al*. 2019). With a wealth of database resources and high-resolution geographic maps, language serves as a valuable comparative tool for studying anthropogenic effects on genetic variation. At a local scale, maize variation has been shown to vary with linguistic differences (Orozco-Ramírez *et al*. 2016; Perales *et al*. 2005), suggesting that trade between communities could be hampered by language barriers. However, the impact of language differences between entire Mesoamerican language families on maize variation is lesser known – little mutual intelligibility exists between a majority of Mesaomerican languages, even for words referring to common trade items such as maize (Barrett 2008; Brown *et al*. 2014). However, extensive long-distance trade could also lead to increased contact between peoples of different ethnolinguistic groups, resulting in the increase of loanwords and mutual intelligibility (Chacon 2022). Thus, both cultural differences across ethnolinguistic groups along with geography could affect maize genetic diversity, making maize an ideal study system to quantify the anthropogenic effects on a domesticate through the lens of bio-cultural coevolution.

Here, we leverage large-scale genotype data of traditional maize varieties (Romero Navarro *et al*. 2017), whole-genome sequencing of human samples from Mexican indigenous populations (Medina-Muñoz 2026), and publicly available linguistic data (Jäger 2018) to measure signals of shared genetic relatedness along geographical and cultural gradients and quantitatively testing the bio-cultural coevolutionary framework. To quantify these relationships, we assess the impacts of geography on maize population structure and generate linear models of maize genotypes to estimate the contributions of human population structure and ethnolinguistic groupings to maize diversity. We find that, in contrast to patterns seen in other species, extant human population structure and ethnolinguistic differences seem to explain a relatively minor amount of modern maize genetic diversity. Finally, to evaluate whether individual loci may show signs of adaptation across ethnolinguistic groups, we perform linguistic GWAS and selection scans to identify loci associated with anthropogenic factors.

## Results

To evaluate the spatial and anthropogenic components of maize genetic diversity, we leverage publicly available sequencing of approximately 2000 traditional varieties of maize (Romero Navarro *et al*. 2017), principal component analysis of whole genome sequencing of 515 human individuals (Medina-Muñoz 2026), and two linguistic datasets that span Mesoamerica and South America (Haynie 2014; Digital 2024). We choose to focus primarily on maize in Mexico, as it is both the center of maize diversity and features the best sampling across these datasets.

### Maize genetic variation is weakly correlated with geography

Using data from a filtered set of 1938 traditional maize varieties (Romero Navarro *et al*. 2017) sampled across Mexico and Central America, we first use FEEMS (Marcus *et al*. 2021) to estimate migration surfaces for maize. Across multiple estimated surfaces, we consistently find decreased gene flow in mountainous regions compared to lowland and coastal regions throughout the country (Fig 1A). To a lesser extent, Oaxaca and the Yucatan Peninsula (Bellon *et al*. 2003; Pressoir and Berthaud 2004) show varying levels of migration when accounting for sampling density bias via downsampling (Fig S1). Overall, these migration surfaces support an important role for elevation in patterning maize diversity, but also suggest the feasibility of long-distance movement across Mexico.

**Figure 1.**
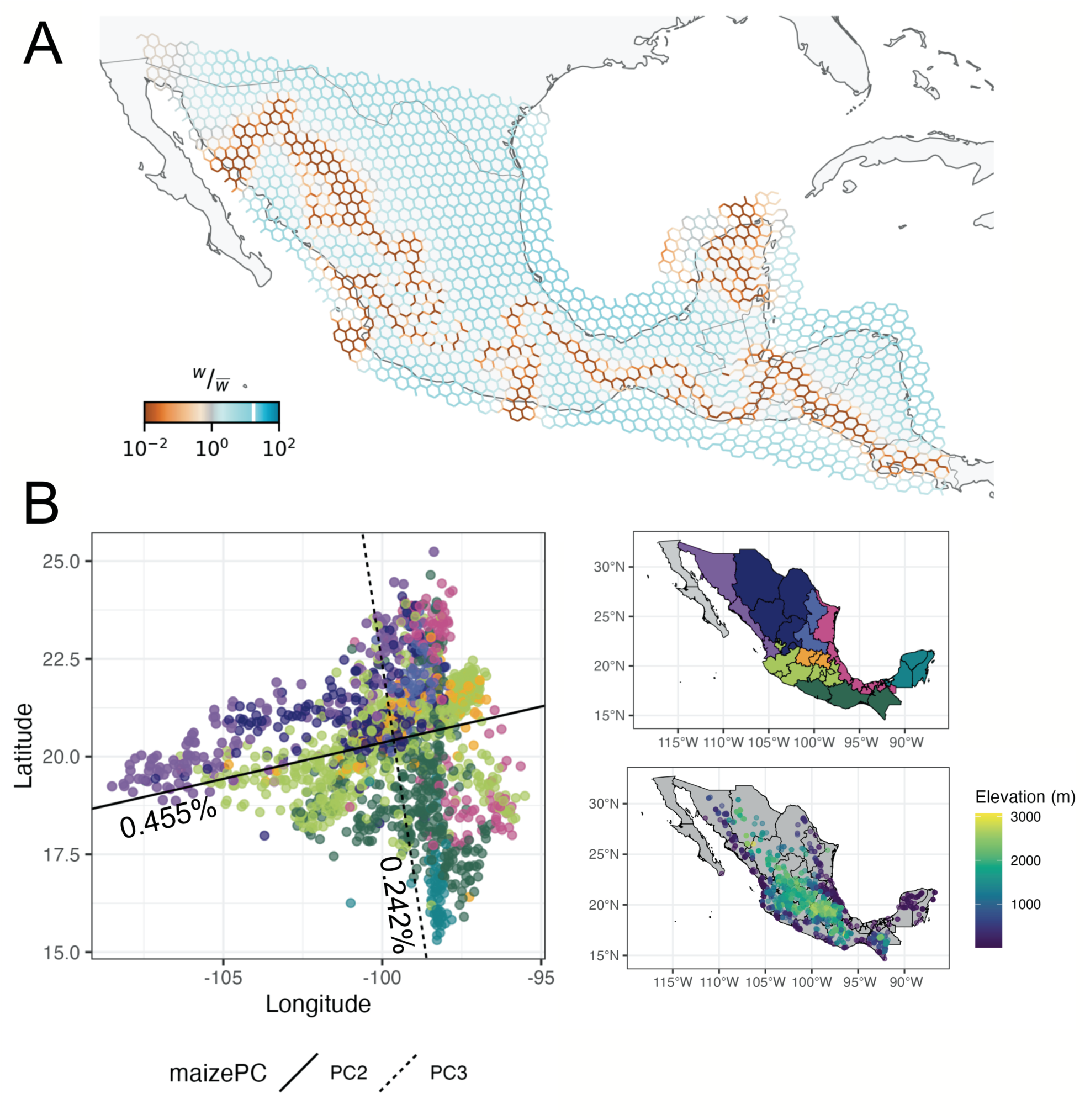
The geography of population structure in maize. **A)** FEEMS plot showing migration surfaces along the Mexico and Central American landscape. Colored edges reflect the estimated migration rate *w* between nodes, with warmer colors indicating higher resistance, and cooler colors indicating higher gene flow. **B)** PC 2 and 3 of Mexican maize samples rotated along latitude and longitude via Procrustes. The percent variance explained by each maize PC is labeled on rotated axes. Points are colored according to their location within nine geographical regions in Mexico: Baja California, Sierra Madre Occidental and Pacific Coastal Lowlands, Mexican Plateau, Sierra Madre Oriental, Gulf Coastal Plain, Mesa Central, Cordillera Neovolcanica, Southern Highlands, Yucatan Peninsula (see upper right map). Elevation of maize sample location indicated by lower right map with warmer colors being higher elevation and cooler colors being lower elevation.

Consistent with the connectivity seen in FEEMS, principal components of maize diversity capture only a fraction of the total variation. PC1, for example, only contributes 1.74%, and the first ten PCs combined explain *<* 5% (Fig S2) As seen in prior genetic work (Yang *et al*. 2023; van Heerwaarden *et al*. 2011), we find that PC1 is highly correlated with elevation (*R*^2^ = 0.971) likely due to admixture in the Mexican highlands with the wild relative *Zea mays* ssp. *mexicana* (Yang *et al*. 2023; van Heerwaarden *et al*. 2011). While weaker than elevation, the second and third PCs nonetheless show substantial correlations with latitude and longitude (*r* = 0.54, Fig 1B). Given the dependence of maize on humans for migration and survival, the overall weak population structure along with correlation to geography point to effective human movement and mixture of maize at large geographic scales. Indeed, data from 515 individuals sampled from 40 population groups associated with distinct linguistic regions across Mexico (Medina-Muñoz 2026) show that the first two genetic PCs of human populations are similarly correlated with latitude and longitude (*r* = 0.576), suggesting that the spatial patterns of Mexican maize may mirror the spatial patterning of humans in Mexico.

### Human genetic variation explains little additional maize genetic variation beyond geography

Similar correlations between genetics and geography in both humans and maize could reflect an anthropogenic-specific effect such as cultural differentiation or selection on maize genetic diversity or simply shared geographic dispersal. To disentangle these possibilities, we pair individual maize collections with nearby human populations (*≤* 21km, but see Supp. Fig S2) for qualitatively similar results using a radius of 100km). We assign to each maize sample the average eigenvalue of the nearby human samples for each human PC (see **Materials and Methods**). Procrustes correlation on this reduced dataset (human *N* = 345, maize *N* = 421) show strong correlations (*r* = 0.43, Fig 2A). Pairwise correlation between all maize PCs and the first 10 human PCs show the strongest correlations with the first three human PCs (Fig 2C). The stronger correlations we detect involve PCs that differentiate North of Mexico or West coast human groups from the rest of Mexico, perhaps due to gaps in the spatial sampling of the human data (Fig S3).

**Figure 2.**
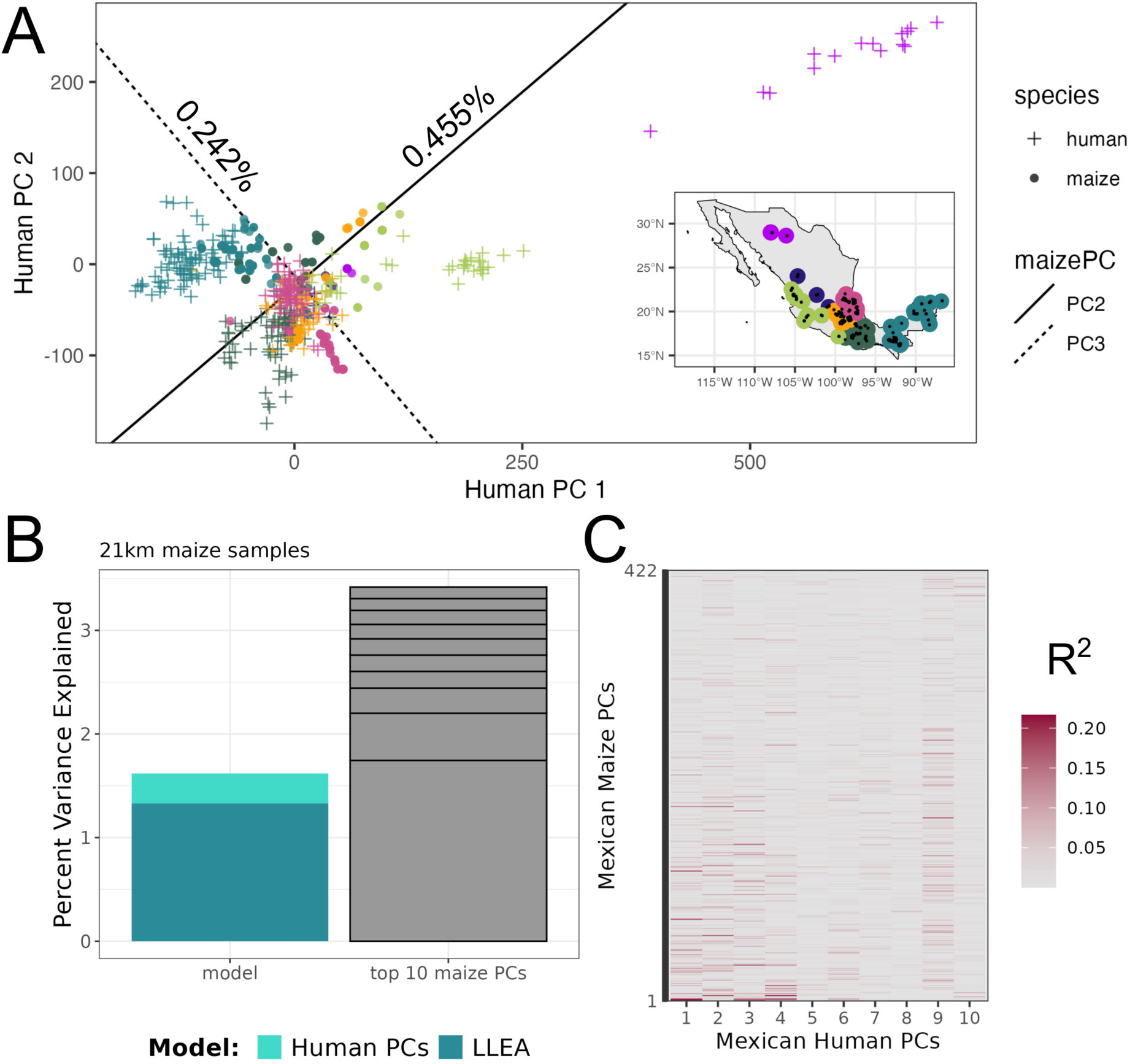
Geography and human population structure have limited impact on maize diversity. **A)** Procrustes of grouped maize (circles) and human (crosses) samples. Inset map indicates geographic position of human samples (colored points) and maize samples (black points). Note that point sizes do not reflect the actual 21 km radius. **B)** Linear model results of maize genetic variation explained by human PCs and geographic features. Genome wide estimated percent maize variance explained by models with just latitude, longitude, elevation, and admixture (LLEA) and the additional amount explained when ten human PCs are included (human PCs). On the right, the first 10 maize PCs are represented as black stacked bars moving from PC 1 to 10 sequentially. **C)** Correlations between maize and human PCs. Darker colors indicate stronger correlations.

To formally quantify the contribution of different processes to maize genetic diversity, we perform a series of linear models for each maize SNP, modeling its variation as caused by a combination of geography, admixture with *mexicana*, and human principal components (Fig 2B; see Methods). The model including geography and admixture explains on average 1.33% of maize genetic variation, and adding the first 10 human PCs captures only an additional 0.29%, less than that explained by the first PC of these maize data (1.745%; Fig 2; Fig S4). Most of the maize genetic variation explained by human PCs in models without geographic parameters is attributed to geographic variables when geography is included (Fig S5). Repeating the models using maize PCs instead of genotypes as the response variable show that human PCs can capture additional variation in the maize PCs beyond geography (Fig S6). However, each maize PC captures less than 1% of the total variation across genotypes, and thus these relationships between maize and human PCs explain little variation within the maize population. Shared geography appears to explain most of the correlation between human and maize PCs, with other factors leading to human genetic structure playing an extremely minor role.

### Linguistic diversity does not strongly pattern maize genetic diversity

As both geography and human population structure explain similar axes of variation in maize diversity, we turned to investigate other methods of anthropogenic influence. Given the very limited role for human population structure in explaining maize diversity, we ask whether interactions between humans explain more, using linguistic relationships as a proxy. Previous work has shown that human genetics, with the exception of Indo-European languages, is only weakly correlated with linguistic differences (Pakendorf 2014; Creanza *et al*. 2015; Barbieri *et al*. 2022; Longobardi *et al*. 2015), motivating our choice to study these separately. Moreover, ethnolinguistic identity has been shown to be an important driver of local genetic and morphological variation in maize (Perales *et al*. 2005; Orozco-Ramírez *et al*. 2016). To quantify the degree to which maize diversity is influenced by ethnolinguistic diversity at a broader scale, we leverage two independent estimates of the geographical extent of indigenous languages (Digital 2024; Haynie and Gavin 2019). We report results using non-overlapping language distributions estimated at the time of European colonization from Haynie and Gavin (2019), but results from the community-sourced Native Land maps (Digital 2024), which allow geographical overlap among languages, are qualitatively similar (Fig S9). We select maize individuals collected within the ranges of three distinct Mesoamerican linguistic families: Uto-Aztecan, Mayan, and Otomanguean (Fig 3A), assigning each individual to a language based on its geographic coordinates.

**Figure 3.**
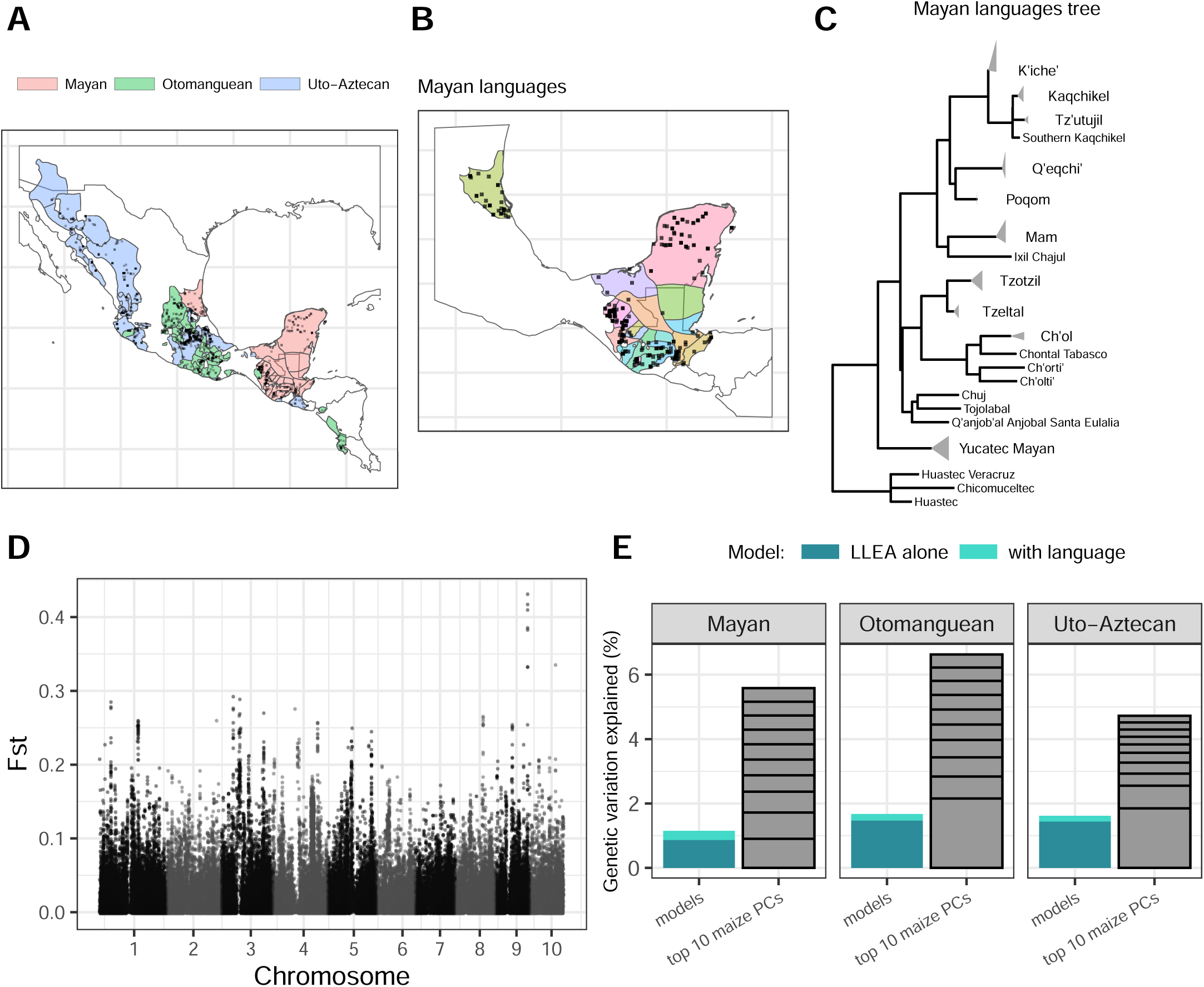
Limited maize genetic variation explained by language diversity. **A)** Language range polygons for Mayan, Otomanguean, and Uto-Aztecan language families in the Haynie dataset, with polygons associated with individual Mayan languages displayed in **B)**. Maize accessions affiliated with languages are shown as points on each map. **C)** Phylogenetic tree of the Mayan language family, with gray triangles clustering dialect groupings within sub-families. Detailed labels of **B)** and **C)** are found in supplemental material. **D)** F_ST_ plots between the three language families for the Haynie dataset. **E)** Proportion of genetic variance explained (bars representing mean adjusted *R*^2^ in %) for each language group. A baseline model of latitude, longitude, elevation, and proportion of *mexicana* admixture (LLEA) was compared to a model including LLEA plus language predictor variables.

Uto-Aztecan, Mayan, and Otomanguean are largely mutually unintelligible, such that linguistic evidence has been unable to reconstruct ancestral relationships between the three (Campbell 1997; Kaufman and Justeson 2009). In spite of these large linguistic differences, maize samples from the three families show little differentiation, with a genome-wide *F_ST_* of only 0.012 (Fig 3B). To evaluate the importance of individual languages, we use language phylogenies from each family (Jäger 2018) to compute the lexical distance between any two languages (Fig 3C and S8). For each maize SNP we run a linear model regression similar to our analysis with human PCs, comparing a base model with geography and admixture (LLEA) to one that includes the first five principal components of the lexical distance matrix (LLEA + language). We find that LLEA models for each of the three families explain similar amounts of genetic variation attributed as in the human PC comparison (1.44%, 0.869%, and 1.47% of maize genetic variation for Uto-Aztecan, Mayan, and Otomanguean families, respectively). Similar to analyses with human PCs, we also find little additional genetic variation attributable to language, with lexical distance explaining 0.18%, 0.29%, and 0.20% additional variation for Uto-Aztecan, Maya, and Otomanguean families, respectively (p-value *<* 10*^−^*^50^ for all models, Fig 3D).

### Linguistic GWAS

While indigenous lexical diversity explains little overall maize genetic variation, patterns of *F_ST_* point to substantial variation along the genome (Fig 3B). For example, among language families, *F_ST_*results identify strong differentiation at a locus on chromosome 9 previously discovered to be associated with climatic variables (Li *et al*. 2025). Genome-wide association (GWAS) of language has previously been used to identify loci potentially under selection in individual cultural groups (Abrouk *et al*. 2020), and we apply this approach to our data, performing case-control GWAS for each language family. By using language as a proxy for anthropogenic factors linked to ethnolinguistic identity, here we identify candidate loci in maize that may be under selection by human cultural practices or adaptations to environment. We restrict our analysis to the Haynie et al (Haynie and Gavin 2019) data set so that each maize individual was attributed to only one language, and grouped similar languages into sub-families to obtain appropriate sample sizes for GWAS (see **Materials and Methods**).

After correcting for multiple testing, we identify 14 putative loci showing significant associations with language. We identify peaks on chromosomes 1, 2, and 8 for a Mayan sub-family consisting of Yucatec, Mopan, and Itza’ Mayan (Fig 4A). We map the distribution of representative SNPs in this peak, showing that the minor allele from the GWAS-identified loci clusters predominantly on the Yucatan peninsula (Fig 4C). In Mayan languages, we additionally identify two putative peaks on chromosomes 5 and 7 in a sub-group of Mayan Ch’ol, Chontal, Ch’orti’, and Ch’olti’ (Fig S10). In Otomanguean languages, we find three peaks on chromosomes 4, 5, and 9 that are highly associated to a grouping of Otomí languages (Fig S10), two peaks on chromosome 8 in Matlatzinca and Mazahua (Fig S10); and a peak on chromosomes 1 and 3 in a sub-group of Mangue, Chiapenec, and Mephaa (Fig S10). Finally, in Uto-Aztecan languages, we also find a significant peak in a sub-family of Cora/Huichol samples in the Uto-Aztecan family (Fig S10) as well as one peak on chromosome 9 in a sub-group of Pipil and closely-related Nahuatl dialects (Fig S10).

**Figure 4.**
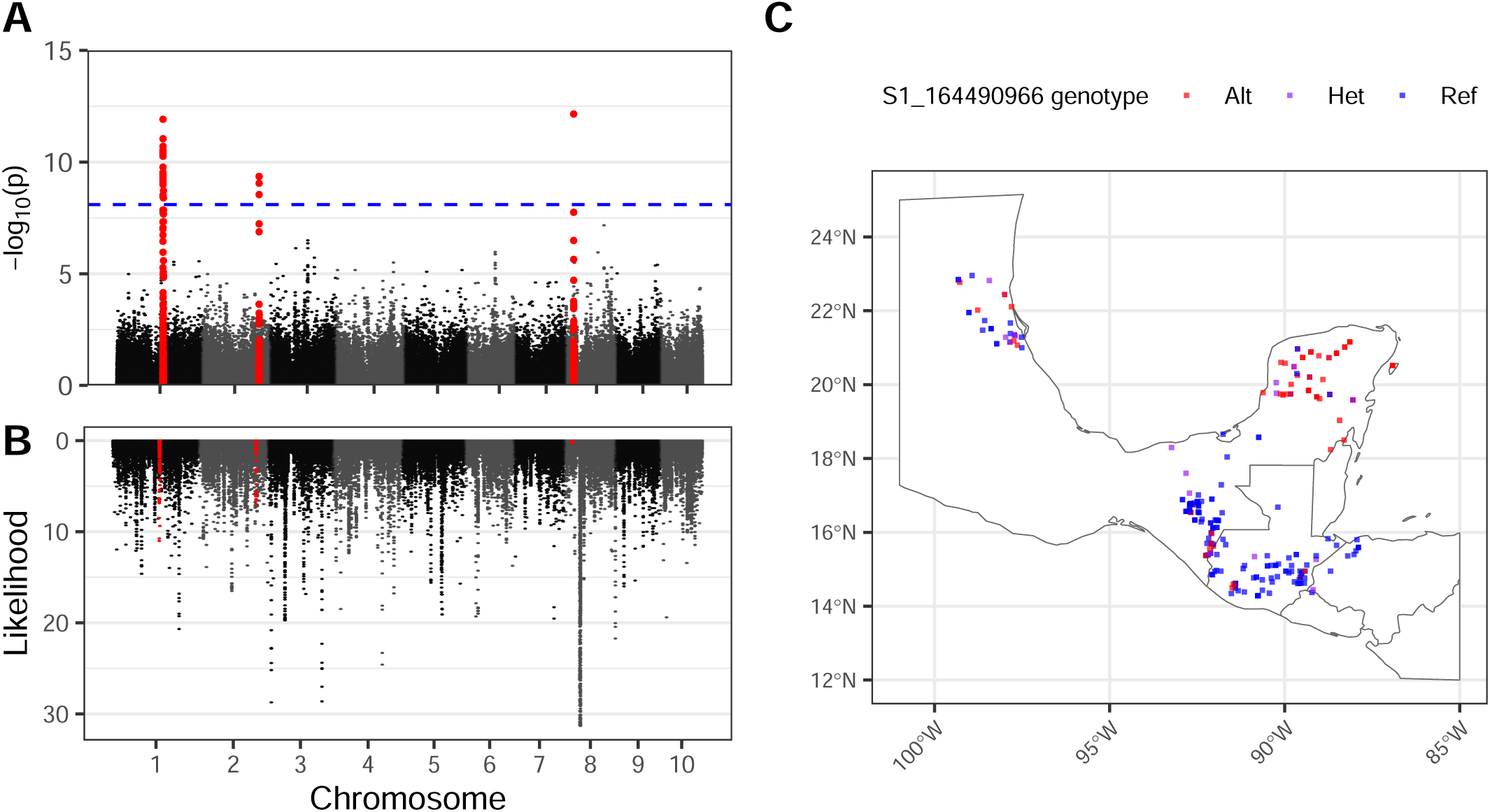
Loci associated with Yucatec Mayan languages show evidence of recent selection. **A)** GWAS shows association with languages in the Yucatec Mayan sub-family including Yucatec Maya, Mopan Maya, and Itza’ Maya. **B)** SFS-based SweeD selection indicates selection within Yucatec Mayan-affiliated maize accessions on chromosomes 1 and 2. **C)** The lead SNP associated with the chromosome 1 GWAS peak shows enriched geographic segregation of the alternate allele in the Yucatan Peninsula. Bonferroni-corrected threshold set to 8e^-9^.

If the associations we find from the linguistic GWAS are in fact due to selection in local ethnolinguistic groups, we would expect to see population genetic signals of selection at these loci. To test this, we performed an SFS-based selection scan using SweeD (Pavlidis *et al*. 2013) for each chromosome exhibiting an association with language. We find evidence consistent with recent selection for the Yucatec Mayan peaks on chromosomes 1 and 2 (Fig 4C), falling within the top 0.07% and 0.30% of composite likelihood ratio (CLR) odds of their respective chromosomes. We additionally identify signals of selection for the peaks on chromosomes 4 and 5 for the Otomí languages that correspond to the linguistic GWAS peak (Fig S11), with CLR scores within the top 0.3% and 0.6% of positions of their respective chromosomes. For the GWAS peaks found in Cora/Huichol, Matlatzinca/Mazhua, Pipil and Nahuatl, Ch’ol/Chontal/Ch’orti’/Ch’olti’, and chromosome 3 for Mangue/Chiapenec/Mephaa, we identify signals of selection, but positions in these loci returned CLR odds from 1% to 10% (Fig S11). We did not find high CLR scores (rank *<* 20th percentile) for the loci in Yucatec Mayan chromosome 8, Otomí chromosome 9, or Mangue/Chiapenec/Mephaa chromosome 9.

A closer look at regions showing evidence of association and selection provides initial suggestions as to the possible functional role for these regions. The locus on chromosome 4 under selection in Otomí, for example, corresponds to a known inversion *Inv4m* that has been linked to flowering time and highland adaptation (Rodríguez-Zapata *et al*. 2026). While associated markers at other loci span relatively large regions (200kb - 2Mb) that include several genes, several genes nonetheless stand out for their potential function. Genes associated with the Yucatec Maya language on chromosome 1 include *xlg1* (Zm00001eb030570), encoding for an extra-large GTP-binding (XLG) protein implicated in abiotic stress (Ding *et al*. 2008; Cantos *et al*. 2023; Wu *et al*. 2018) and *umc2232* (Zm00001eb030420) which has been characterized as a calcium-transporting ATPase, suggesting a potential role in adaptation to limestone-rich soils in the Yucatan peninsula. And on chromosome 2, associations with Yucatec Maya include genes involved in vitamin B synthesis (*qpt1*; Zm00001eb104420 (Suzuki *et al*. 2020)) and flavor and sugar storage in waxy and sweet corn (*ugp1*; Zm00001eb104400 (Luo *et al*. 2024)), suggesting possible selection for culinary preference or nutritional quality.

## Discussion

Identifying and quantifying anthropogenic factors behind the diversity of domesticated crops remains a challenging affair due to the myriad of correlated factors involved in their evolution. Using sequencing and passport data from approximately 2,000 traditional maize varieties and 515 human genomes representing samples from across Mexico, we quantify the impacts of geography and human genetic and cultural diversity on genetic variation in maize.

We find that maize genetic diversity is largely unstructured, with no single feature explaining more than a few percent of the variation and the top 10 principal components accounting for less than 5% of the total genetic variation. FEEMS spatial analysis reveals that gene flow is far from homogeneous across the landscape, with reduced movement in the highlands, but range-wide *F_ST_* results nonetheless show that, overall, such barriers have not led to the accumulation of strong genetic differentiation. Maize genetic PCs correlate strongly with both geography and human population structure, likely driven by the dependence of maize on human dispersal and the shared geographic history of the species. However, each maize PC captures little of the total genetic variation and thus geography and human population structure are likely weaker drivers of total maize genetic diversity. Similarly, we show that linguistic diversity among indigenous languages provides only marginal explanatory power for large-scale patterns of maize diversity, explaining approximately one percent of genetic variation.

Our finding stands in contrast to previous work showing that incorporation of language variables captured meaningful genetic differentiation in crops such as cassava Delêtre *et al*. (2021), sorghum Labeyrie *et al*. (2014, 2016), pearl millet Naino Jika *et al*. (2017), and fonio millet Abrouk *et al*. (2020); Diop *et al*. (2025). One major difference between our work and these other studies is that we investigate a broader geographic scale (but see Gilabert *et al*. 2025; Cerda *et al*. 2025). This difference in scale not only increases the total genetic diversity to be explained, but also diminishes the role of heterogeneous local processes that may be important at smaller, local scales (Sakamoto *et al*. 2024; Lasky *et al*. 2024). In one study of pearl millet, for example, a single lake explains a meaningful proportion of the observed population structure (Naino Jika *et al*. 2017). This difference in scale and process may also help explain why our results differ from a previous maize analysis that focused on two neighboring municipalities in Oaxaca, in which ethnolinguistic group was confounded with the presence of roads between villages, likely facilitating trade between villages of the same ethnolinguistic group (Orozco-Ramírez *et al*. 2016). Indeed, large-scale drivers of environmental variation that we see correlate broadly with human and maize variation across Mexico play less of a role in investigations of other crops at smaller spatial scales (e.g. Labeyrie *et al*. 2014, 2016). Of course, there are also numerous biological and cultural differences between maize and other studied crops which could explain the different patterns observed.

In maize, a long history of extensive trade could have prevented the accumulation of genetic differentiation among populations. Archaeological evidence demonstrates frequent long-distance trade occurred throughout Mexico and Mesoamerica (Hirth 1978; Chase *et al*. 2025; Golden *et al*. 2012; Palka 2024). These trade routes were co-opted by Spanish colonizers and persist in regional markets to the present day (Linares and Bye 2016), and maize farmers continue to maintain extensive networks exchanging both seed and cultivation practices (Bellon *et al*. 2018). Efficient trade networks have also been facilitated by widespread use of common language for trade. Historically, Mayan and Nahuatl dialects acted as *lingua franca* at the height of a multilingual Aztecan empire (Braarvig and Geller 2018; Dakin 2010; Peckham 2012), and in post-colonial Mexico and Central America, Spanish has played an analogous role in shaping communication, trade, and migration.

While genome-wide diversity shows little correlation with within-language family diversity, we nonetheless find individual genetic variants associated with linguistic subfamilies. Regions identified by GWAS are wide and encompass many putative genes, but several include genes consistent with anthropogenic or abiotic selection. For instance, loci associated with the Yucatec Maya language include *ugp1*, previously linked to sugar deposition in kernels that may affect both taste and storage longevity (Luo *et al*. 2024), and the genes (*xlg1* and *umc2232*), which may have been selected by agricultural practices tailored to the calcium-rich and alkaline soil (Voorhies and Michaels 2024). Similarly, the inversion polymorphism *Inv4m* that we find associated with Otomanguean Otomí languages has been linked to flowering time (Rodríguez-Zapata *et al*. 2026) and identified as important for highland adaptation (Romero Navarro *et al*. 2017; Li *et al*. 2025). In other cases, associations with Otomí or Yucatec Maya languages show evidence of selection without clear candidate loci, and many of the associated loci show no signs of recent selection. Some of these may be false positives due to confounding with population structure or spatial auto-correlation, but others may reflect more complex forms of selection. Further fine-mapping within these GWAS intervals may be useful in identifying potential causes of selection.

Moving forward, our work provides a quantitative framework for investigation of bio-cultural coevolution in anthropogenic systems and highlights the many factors that may impact this process. An important avenue for future efforts will be improving our understanding of historical movement patterns for both humans and their domesticates. Recent work in both dogs (Bergström *et al*. 2020a; Zhang *et al*. 2025) and sheep (Daly *et al*. 2025), for example, use ancient DNA to highlight how patterns of spatial diversity in domesticates reflect known human dispersal paths. As in maize, however, trade and cultural preferences may distort these simple patterns – long distance trade likely explains conflicting patterns of diversity between dogs and humans in some regions (Bergström *et al*. 2020a; Zhang *et al*. 2025), and both introgression and cultural preferences impacted sheep diversity during Bronze Age migration from Asia into Europe (Daly *et al*. 2025). While there are clearly many complexities to these studies, combined they highlight the power of dense, multi-scale geographic sampling and the utility of genome-wide data, including ancient DNA, to understand patterns and principles of bio-cultural coevolution.

## Materials and Methods

### Maize Sampling and Genotyping

Maize accessions were sampled from the CIMMYT seed bank Seeds of Discovery (SeeDs) dataset (see (Romero Navarro *et al*. 2017) for a more detailed description of this SNP dataset). Of 4693 initial accessions with genotyped data, we first limited to 3511 accessions with available geo-coordinate data.

For FEEMS migration surface analysis, we used a subset of 1939 maize accessions from Mexico, Guatemala, Belize, El Salvador, Honduras, Nicaragua, and Costa Rica. For human-maize Procrustes and regression analyses, maize samples were subset to those which occurred within 21 km (human *N* = 345, maize *N* = 421) and 100km (human *N* = 523, maize *N* = 1437) of a human sample using QGIS 3.36.2-Maidenhead QGIS Association (2025) and *r-geosphere* v1.5-18 package (Robert J. Hijmans 2024). Since each of these subsets only contained samples from Mexico, the principal components were calculated using only samples from Mexico for both species (human *N* = 515, maize *N* = 1725). For linguistic analyses, a spatial join was used to find maize accessions whose collection coordinates overlapped with each set of language range polygons (see **Linguistic data**) using the *r-spatial* package (Ripley 2009). For the Native Land polygon ranges, we kept 649, 302, and 320 accessions for the Uto-Aztecan, Mayan, and Otomanguean language families, respectively. For the Haynie polygon ranges, we kept 729, 299, and 295 accessions for the Uto-Aztecan, Mayan, and Otomanguean language families, respectively.

Genotyping-by-sequencing data was acquired from (Romero Navarro *et al*. 2017) and processed as per (Li *et al*. 2025) for an initial set of 856,300 SNPs. For each accession, we used the estimated proportion of admixture with the wild relative *Zea mays* ssp. *mexicana* estimated by Yang *et al*. (2023).

### Human and maize PCA Procrustes

SNPs with a minor allele frequency less than 0.01 were removed with bcftools v1.19 (Danecek *et al*. 2021) and LD-pruned (*r*^2^ *>* 0.1) with PLINK v2 (Chang *et al*. 2015). Missing genotypes were imputed with the mean genotype. Maize genotypes were regressed against admixture proportion and then residuals were used as admixture removed genotypes. PCs 2 and 3 calculated from the non-transformed maize genotypes were also used in Procrustes analyses and generated qualitatively similar results as the admixture removed maize genotypes (Fig S6). For clarity and ease, we present here analyses that use non-transformed maize genotypes instead of the admixture removed genotypes. PCs were calculated in R 4.4.2 using ‘prcomp’ (R Core Team 2024).

We also used the first 10 PCs calculated from admixture-removed whole-genome sequence of 515 Mexican indigenous individuals from Mexican Biobank and the GenomeAsia 100K project (for more detailed description of this SNP data, see (Gusareva *et al*. 2025; Sohail *et al*. 2023; Moreno-Estrada *et al*. 2014; Bergström *et al*. 2020b)), downloaded from (Medina-Muñoz 2026). Procrustes analyses were calculated using the *vegan* R package 2.6-10 (Jari Oksanen *et al*. 2025) either between samples’ genetic PCs and geography or between the genetic PCs of the two species. To pair maize and human samples, each maize sample was assigned humans within either a 21 or 100 km radius. For each maize sample, the eigenvalues of all assigned human samples were averaged, creating a centroid position for the human samples associated to a single maize individual in human PC space.

### Migration surfaces

We used Fast Effective Estimation of Migration Surfaces (FEEMS) (Marcus *et al*. 2021) to estimate migration rates along the landscape for accessions located within Mesoamerican countries (see above). A discrete global grid at the 50km resolution was used from Sahr *et al*. (2003), with convex boundaries for the local geography of interest drawn manually using https://www.keene.edu/campus/maps/tool/.

FEEMS input took in latitude/longitude coordinates for all samples, as well as SNP genotypes filtered to MAF *>* 0.01 and LD pruned (*r*^2^ = 0.3 within 10kb), for a total of 1939 samples and 232,225 SNP markers. Because we found that edge weights were influenced by sampling density, we tested to see if uniformly sampled nodes resulted in differences in the migration surface patterns. We downsampled to only 1 sample per node and ran the default FEEMS and a newer version that fits node-specific variances (Fig S1) after performing cross-validation for parameterization. The FEEMS weight graph without downsampling was parameterized with a *λ* smoothness turning parameter value of 2, and a *λ*_q_ parameter for node-specific variance penalization of 100. For our downsampled FEEMS graph, we parameterized with *λ* = 12 and *λ*_q_ = 0.01.

### Linguistic data

To quantify similarity between languages, we first gathered languages that had lexical distances computed for the Automated Similarity Judgment Program (ASJP) from Jäger (2018). Briefly, to estimate these language distances, 40-item core vocabulary lists from the ASJP database were used to calculate pointwise mutual information between sound classes. A normalized Levenshtein distance was then calculated from the pointwise mutual information scores to generate a distance matrix and phylogenetic relationship among the languages.

Linguistic range maps were obtained from the Native Land Digital community resource (Digital 2024) and from a dataset associated with Haynie and Gavin (2019) (hereafter referred to as the NL and Haynie datasets, respectively). NL map polygons are crowd-sourced and curated from community resources. Haynie ranges attempt to establish pre-Columbian contact ranges for each language that was able to be matched to a language-level Glottolog identifier (Nordhoff and Hammarström 2011). Briefly, maps sourced for the Haynie ranges were prioritized for those closest to point of European contact, based on previous linguistic research, spatial resolution, and aggregation across multiple sources, and computationally predicted into output polygons. NL range maps encompassed overlapping polygons due to the community curation nature of its origin, whereas Haynie language maps are non-overlapping, meaning that any given maize collection was associated with only one language. For each of these two range datasets, three language families were chosen for their prevalence in Mesoamerica - Uto-Aztecan, Mayan, and Otomanguean. As not all languages present in the NL or Haynie polygons had perfect matches to the ASJP dataset, best efforts were made to manually match language polygon names and ranges to an identical or corresponding language in the ASJP set. Additional details can be found in Supplemental Methods, and the final pairings used in analysis can be found in Supp Data S2.

To calculate lexical distance between two maize accessions, we compared two types of distance metrics. The lexical distance was computed as the Levenshtein distance between the ASJP languages, mapped from the map polygon languages, attributed to each maize accession. For attributed languages that were mapped to multiple ASJP languages or dialects, the lexical distance was instead computed as the average Levenshtein distance between the mapped ASJP language(s) of one accession to all the ASJP language(s) of the other accession. If a maize accession lay within the boundaries of two or more polygons, we also averaged the Levenshtein and branch length distance between each attributed polygon language, on top of the mapped ASJP languages. This helped to account for situations in which a given maize seed lot may be cultivated or traded by a farmer speaking any combination of the languages of the associated polygons.

### Diversity characterization

F_ST_ statistics were calculated using VCFtools v4.2 (Danecek *et al*. 2011). For both the NL and Haynie datasets, accessions attributed to a given language family (Uto-Aztecan, Mayan, and Otomanguean) were each considered as a population. The Weir and Cockerham F_ST_ statistic was calculated between the three populations for each SNP, and the mean F_ST_ across all SNPs was reported.

### Variance partitioning

To compute the amount of genetic variance in the maize population explained by human PCs, language, or other landscape variables, we ran a multivariate regression of genotype on various predictor variables. For both the human PC and language variance partitioning experiments, we compared two hypotheses: a model containing underlying landscape variables that affect genetics, including latitude, longitude, elevation, and proportion of *mexicana* admixture (LLEA), and a model including the LLEA variables plus additional variables for either human PCs or lexical distance (henceforth ANTH). We constructed our LLEA models using point values directly from locations-of-origin. Latitude and longitude (in degrees), and elevation in meters above sea level (masl) variables for each accession were directly taken from CIMMYT passport data.

For the LLEA + human PCs model, we used centroids of the human eigenvectors assigned to each maize sample to reduce many-to-many to one-to-one matching. For the LLEA + language model, as there are no point values for language corresponding to a given accession, we performed a principal coordinates analysis (PCoA) of the pairwise lexical distance matrix using the ‘wcmdscale’ function from the *vegan* package (Jari Oksanen *et al*. 2025), keeping the top *k* = 5 PCoA loading values for each accession. For each linguistic dataset (language + NL/Haynie combination), we used PLINK 1.9 (Chang *et al*. 2015) to filter down the GBS genotype data to a minimum minor allele frequency (MAF) *>* 0.01, and used the *–indep-pairwise* argument to prune SNPs within 50 variants within LD above *r*^2^ *>* 0.1, resulting in a set of 130-140k biallelic SNPs in each dataset. We then converted our SNP genotypes into 0/1/2 dosages for each accession in the dataset. We regressed a vector *y* of SNP genotypes for each accession *j* against the predictor variable terms (a total of 4 predictors for the LLEA model, 14 predictors for the LLEA + human PCs model, and 9 predictors for the LLEA + language model).

The general model was as follows:

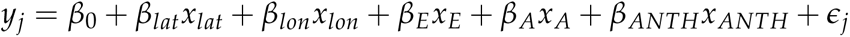

for *j* samples and repeated for *i* SNP markers, and where *β*_0_ is the intercept of the regression. *x_lat_ x_lon_*, *x_E_* and *x_A_* correspond to the *j ×* 1 vectors of latitude, longitude, elevation, and admixture, respectively. *x_ANTH_* represents a general expansion of either the ten terms representing the 10 *j ×* 1 vectors corresponding to the top 10 PCA loadings of the human genetic data in the human regression model, or the 5 *j ×* 1 vectors corresponding to the top five PCoA loadings from the lexical distance matrix for the language regression model. *ɛ* is error, or the relationship between the genotype dosage and unmeasured or cryptic factors. Within these models, we do not pull out individual SNPs as associated with ANTH variables but rather average across all SNPs to get a genome-wide estimate of the variance captured by each set of parameters.

To find the proportion of genetic variance *R*^2^ explained by human PCs or language, we ran the LLEA model without the *β_ANTH_ x_ANTH_* terms and compared it to the equivalent models containing both the LLEA and ANTH terms. Using a linear model with the *lm* function, we computed the *R*^2^ value for each SNP for both models. To account for the different number of predictors in the LLEA and LLEA + ANTH models, we used the adjusted *R*^2^ value reported. We subtracted the adjusted mean *R*^2^ from each LLEA only model from the adjusted *R*^2^ of the corresponding LLEA + ANTH model for each SNP and took the mean to obtain the genetic variation explained by only ANTH:

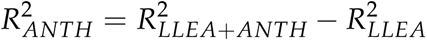

### Linguistic GWAS

To evaluate associations between genetic variants and ethnolinguistic identity, case-control GWAS was run on larger groupings of languages. As sample sizes of accessions were too small for GWAS analysis for individual languages, we grouped multiple languages together into 21 sub-groupings based on phylogenetic relationships in the ASJP similarity dataset. Languages were grouped together based on phylogenetic common ancestor until most sub-groupings reached at least n = 15. Notably, Chinantecan, a language with many highly mutually unintelligible dialects, was not easily grouped with other language sub-groupings, resulting in only 5 accessions, and was thus excluded from GWAS. The Eastern Popolocan/Western Popolocan/Chochotec as well as the Chuj/Tojolabl/Q’anjob’al sub-groupings also had low sample sizes (less than 20) and were not included for GWAS. Full groupings can be found at Supp Data S1.

GEMMA was used to perform case-control GWAS (Zhou and Stephens 2012) for each language grouping against all other accessions in the language family. For this analysis, we only used accessions affiliated with language polygons from the Haynie dataset, such that each accession would only be counted once over all the GWAS. GWAS was performed with GEMMA default parameters, filtering out genotypes with missingness *>* 0.05 and a minor allele frequency *>* 0.01. To account for population structure, a kinship matrix for all accessions in the GWAS was included. A Bonferrroni correction to account for number of individual SNP tests and number of individual language tests was included in the p-value threshold of 8 *×* 10*^−^*^9^.

### Selection scan

SweeD v3.1 (Pavlidis *et al*. 2013) was used to perform selection scans on populations and chromosomes that were marked as potential candidates from GWAS. SweeD was run on chromosomes 4, 5 and 9 for individuals attributed to the grouping of Querétaro, Tilapa, Mezauital, and Ixtenco Otomí languages (33 individuals), chromosome 4 for Cora and Huichol (50 individuals), chromosome 5 and 7 for Ch’ol, Chontal, Ch’orti’, and Ch’olti’ (22 individuals), chromosome 1 and 3 for Mangue, Chiapenec, and Mephaa (25 individuals), chromosome 8 for Matlatzinca and Mazahua (25 individuals), chromosome 9 for Pipil, Northern Puebla Nahuatl, and Eastern Nahuatl (167 individuals), and all chromosomes for individuals attributed to the Yucatec, Mopan, and Itza’ Mayan grouping (64 individuals). Site frequency spectra (SFS) were folded and 40000 positions were taken in each chromosomal alignment to generate the composite likelihood ratio of a selective sweep at each position.

## Data availability

Code for human-maize genetic comparisons and Procrustes analyses are available on Github at https://github.com/Snodgras/maize-human-procrustes. Code for linguistic analyses are available at https://github.com/liforrest6/maize-linguistics. Imputed GBS SNP data used for genetic analysis is available at https://data.cimmyt.org/dataset.xhtml?persistentId=hdl:11529/8702394. Human PC data is available at (Medina-Muñoz 2026)

## Acknowledgments

Map data provided by Native Land Digital (https://native-land.ca/), used with permission for educational and non-commercial purposes. We would like to acknowledge David Mora Marin for advice on linguistics, Hannah Haynie for early access to her linguistic map, and James Kitchens for advice and help on GIS and allele frequency surfaces.

## Author Contributions

SS, FL, JRI, GC, DR, AME designed research SS, FL, SM, SMM, LS, CP, MM, NK, AGG, SCS, HLK performed research; SS, FL, SM, SMM, LS, CP analyzed data; JRI, DR, GC supervised the research; and SS, FL, LS, SM, DR, GC, JRI wrote the paper.

## Supplementary Tables and Figures

**Figure S1.**
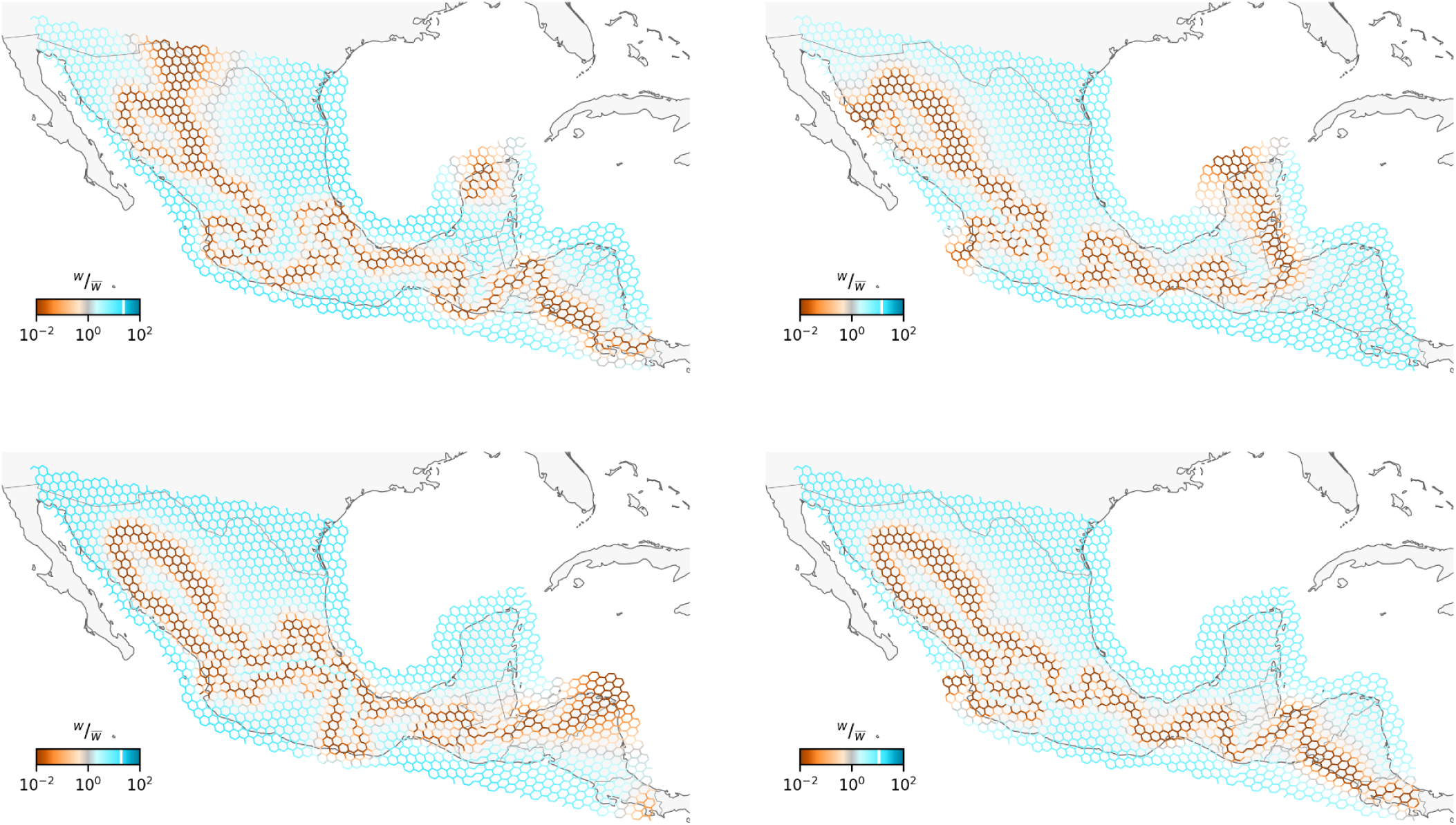
Demonstration of FEEMS down-sampling. Example of four FEEMS runs after down-sampling to maximum one sample per node, with parameters chosen from cross-validation (*λ* = 12, *λ*_q_ = 0.01) with node-specific variance estimation.

**Figure S2.**
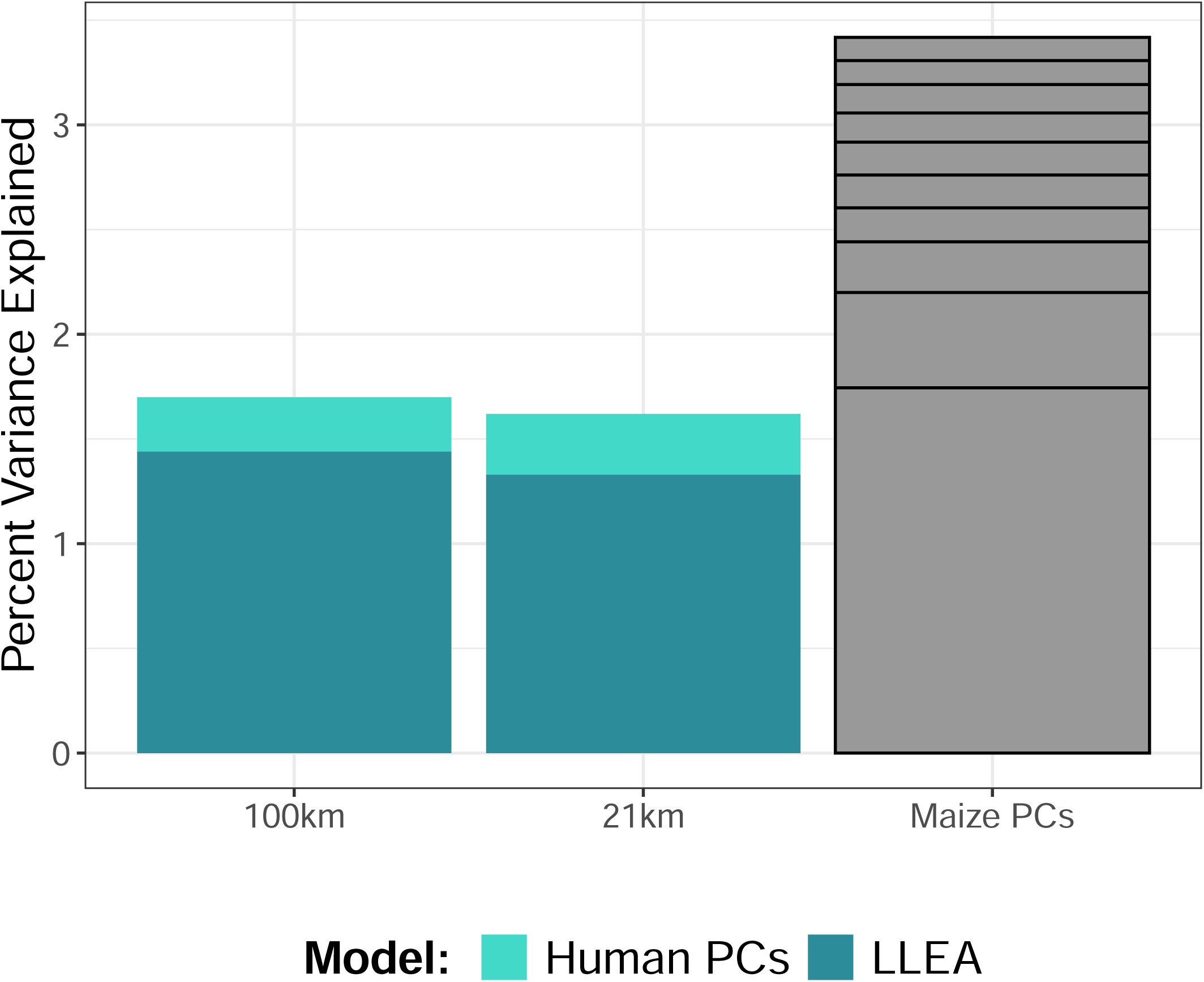
Comparison of 100km vs. 21km pairing models. Humans and maize samples were paired either with 100km or 21km radii. The average percent variance explained of maize genotypes from genome-wide regression analysis are plotted for each pairing distance. Dark blue is the percent variance explained solely using latitude, longitude, elevation, and admixture with *mexicana* proportion (LLEA) as parameters. The light blue indicates the additional percent variance explained when the top ten human PCs are included as parameters. For context, the percent variance explained by each of the top ten maize PCs, starting from PC 1 up through PC 10, are plotted in grey.

**Figure S3.** Correlations between maize and human PCs. The first plot shows the color scheme used to denote the various ethnolingustic regions of the human samples and their location within Mexico. This color scheme is used throughout. All following plots are individual maize and human PC correlations that are greater than 0.6. Lines represent linear regression line of best fit. Titles of plots include the correlation and *R*^2^ values of each set of PCs.

**Figure S4.**
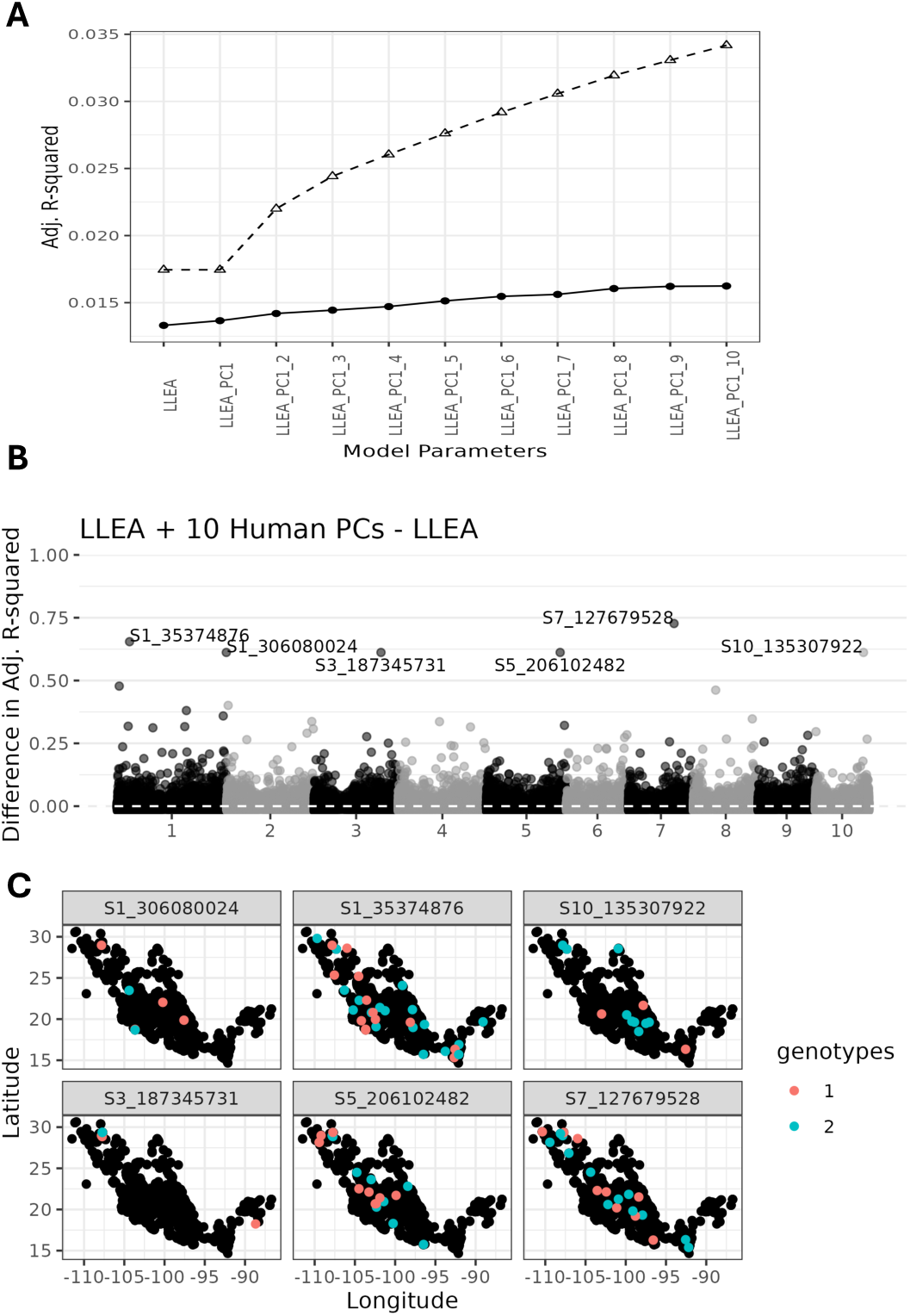
Results of maize and human PC regression analysis. **(A)** The average adjusted *R*^2^ values as human PCs are sequentially added as model parameters. The solid line and circles are the results from the regression models and the dashed lines and triangles are the cumulative sum of the top ten maize PCs. The value of the first maize PC is plotted twice for the LLEA model and the LLEA + human PC1 model. **(B)** The difference between the adjusted *R*^2^ values for the full parameter model and the LLEA only model per SNP is plotted by genomic position. This difference represents the additional *R*^2^ captured with inclusion of the human PCs as model parameters. The white dashed line highlights the 0 value. Six top snps are highlighted as showing a strong association with human parameters. **(C)** Genotypes of each of the top six SNPs identified in B across Mexico. Black points represent samples with the reference allele. Red and blue points represent samples heterozygous or homozygous for the alternative allele respectively.

**Figure S5.**
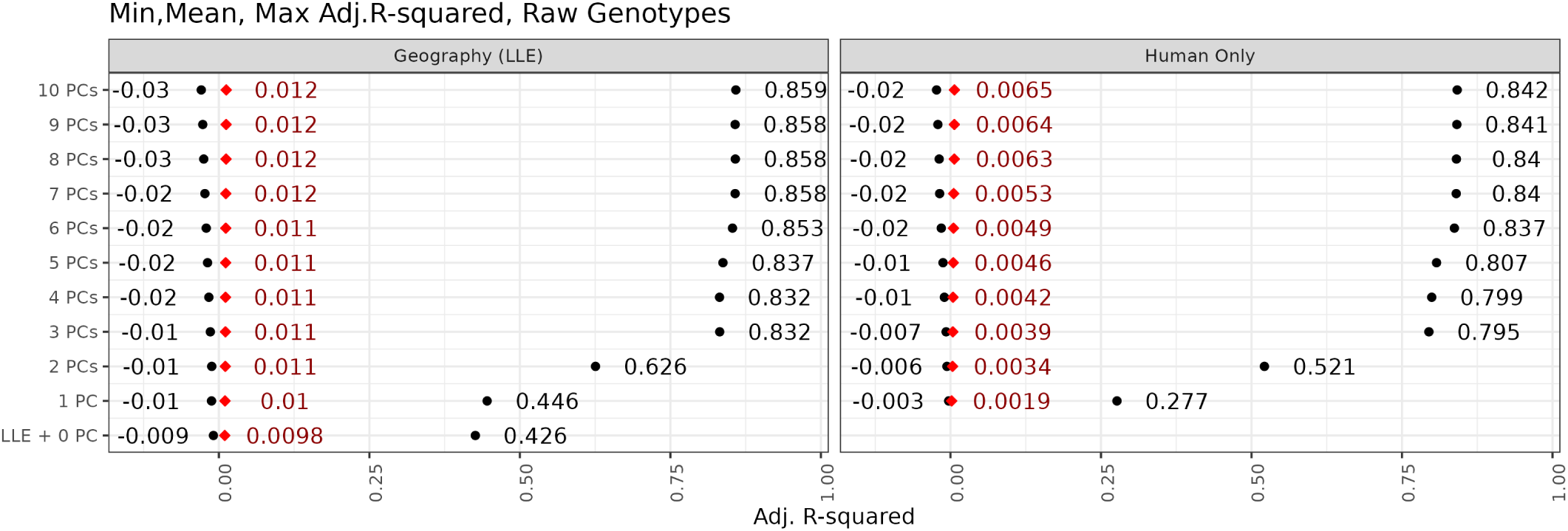
Minimum, mean, and maximum adjusted. *R*^2^ **values for human PC linear regressions** Adjusted *R*^2^ values for linear regression as each human PC is added as a parameter, with the full model being all 10 human PCs. Models in the Geography (LLE) plot also include latitude, longitude, and elevation as model parameters. Values for each point are written to the side.

**Figure S6.**
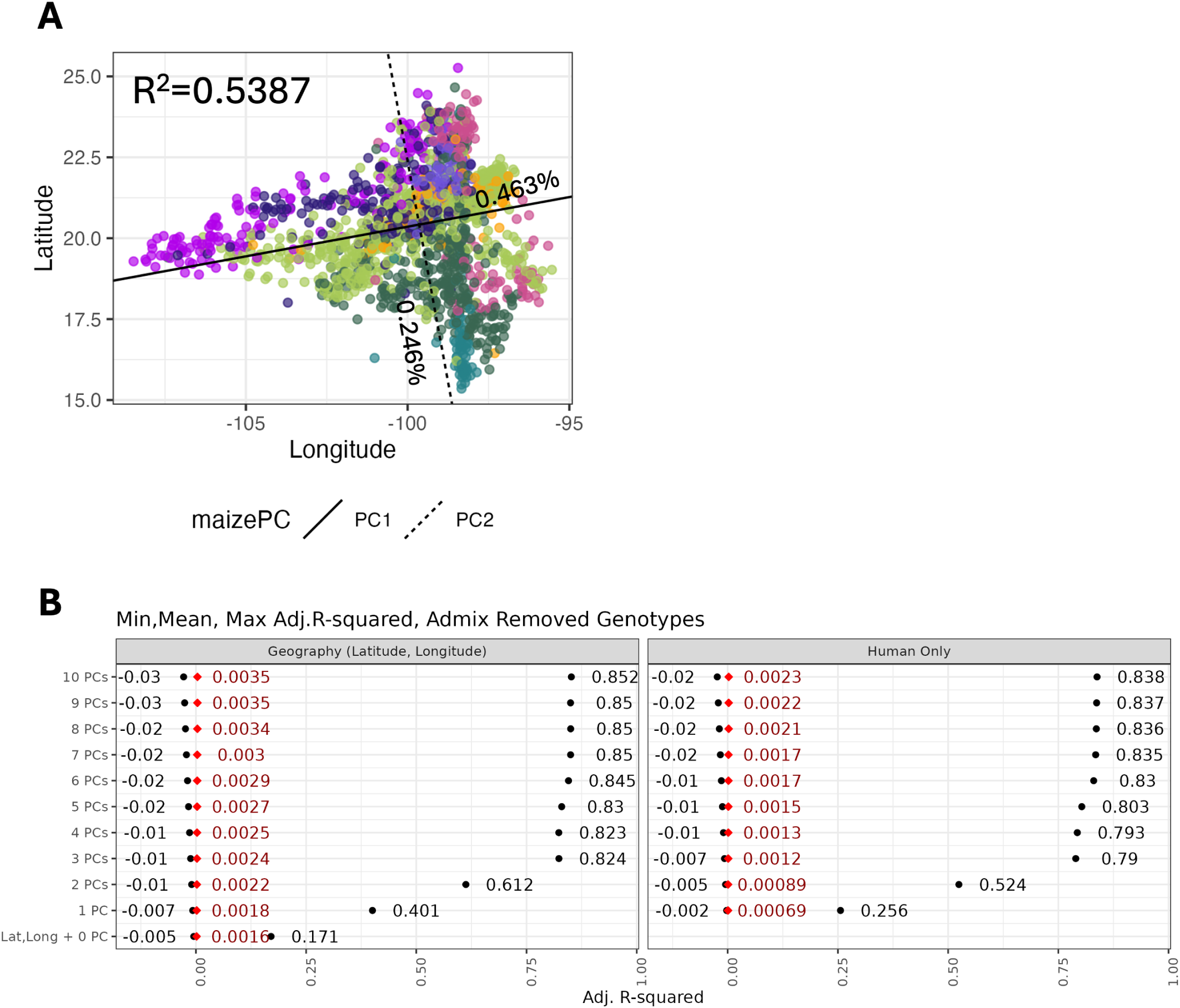
Results from using admixture removed maize genotypes. **(A)** Procrustes rotation of the first two admixture removed maize PCs onto sample geographic coordinates. Results are nearly identical to non-transformed genotypes using PC 2 and 3. **(B)** Linear regression results using admixture removed maize genotypes with human PCs and environmental parameters latitude and longitude. Points represent minimum, mean, and maximum adjusted *R*^2^ across models. Point values are written beside the points.

**Figure S7.**
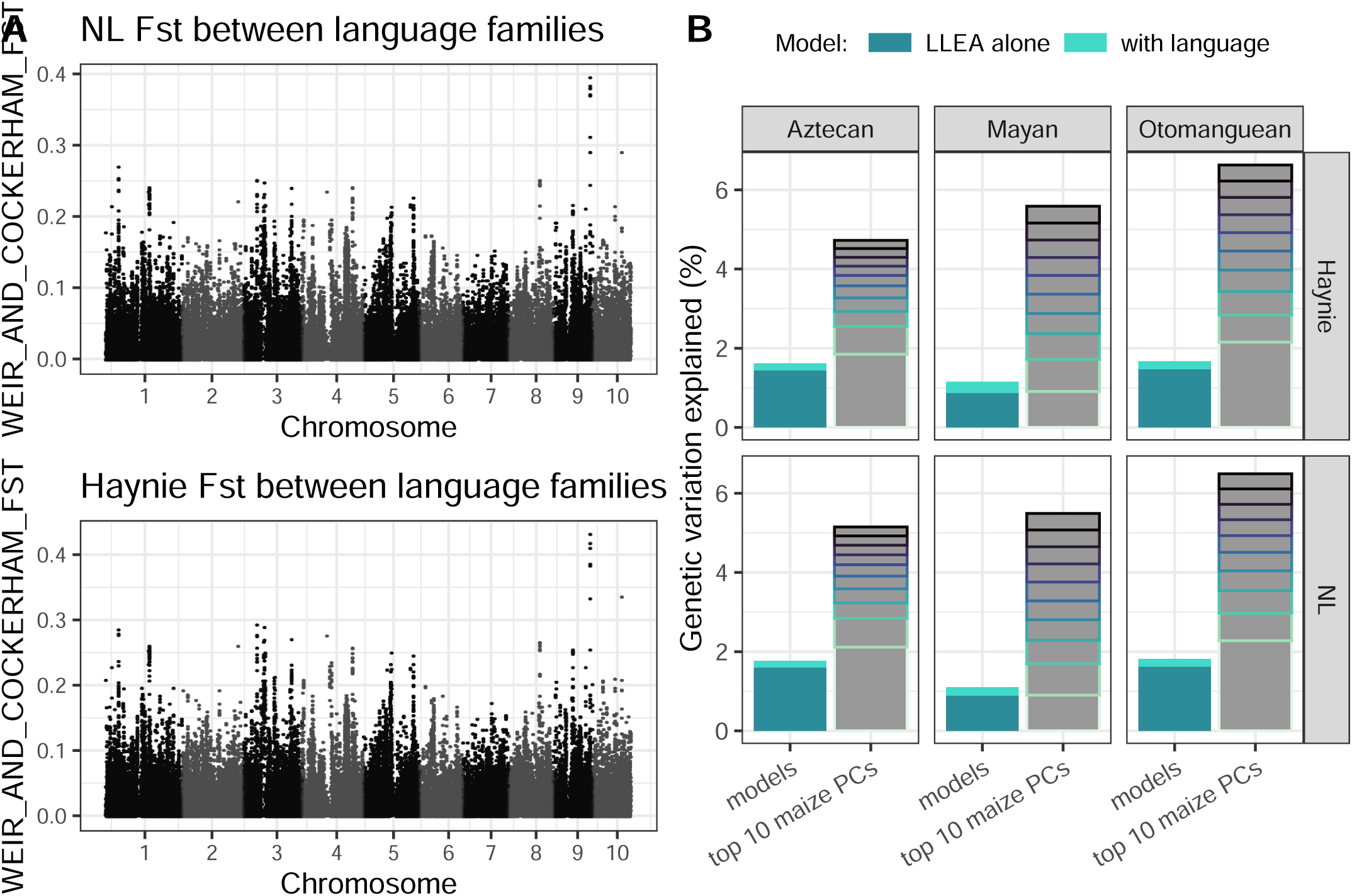
Comparison of Native Land dataset to Haynie. **A)** Comparison of genome-wide *F_ST_* in Haynie dataset as compared to NL, **B)** Comparision of variance partitioning in NL dataset as compared to Haynie

**Figure S8.** All linguistic trees. Sub-groupings used for variance partitioning labeled by color. Individual languages not grouped are colored gray.

**Figure S9.** All language maps. Maps for each language range in each dataset and language. Otomanguean NL, Otomanguean Haynie, Uto-Aztecan NL, Uto-Aztecan Haynie, Mayan NL, Mayan Haynie

**Figure S10.**
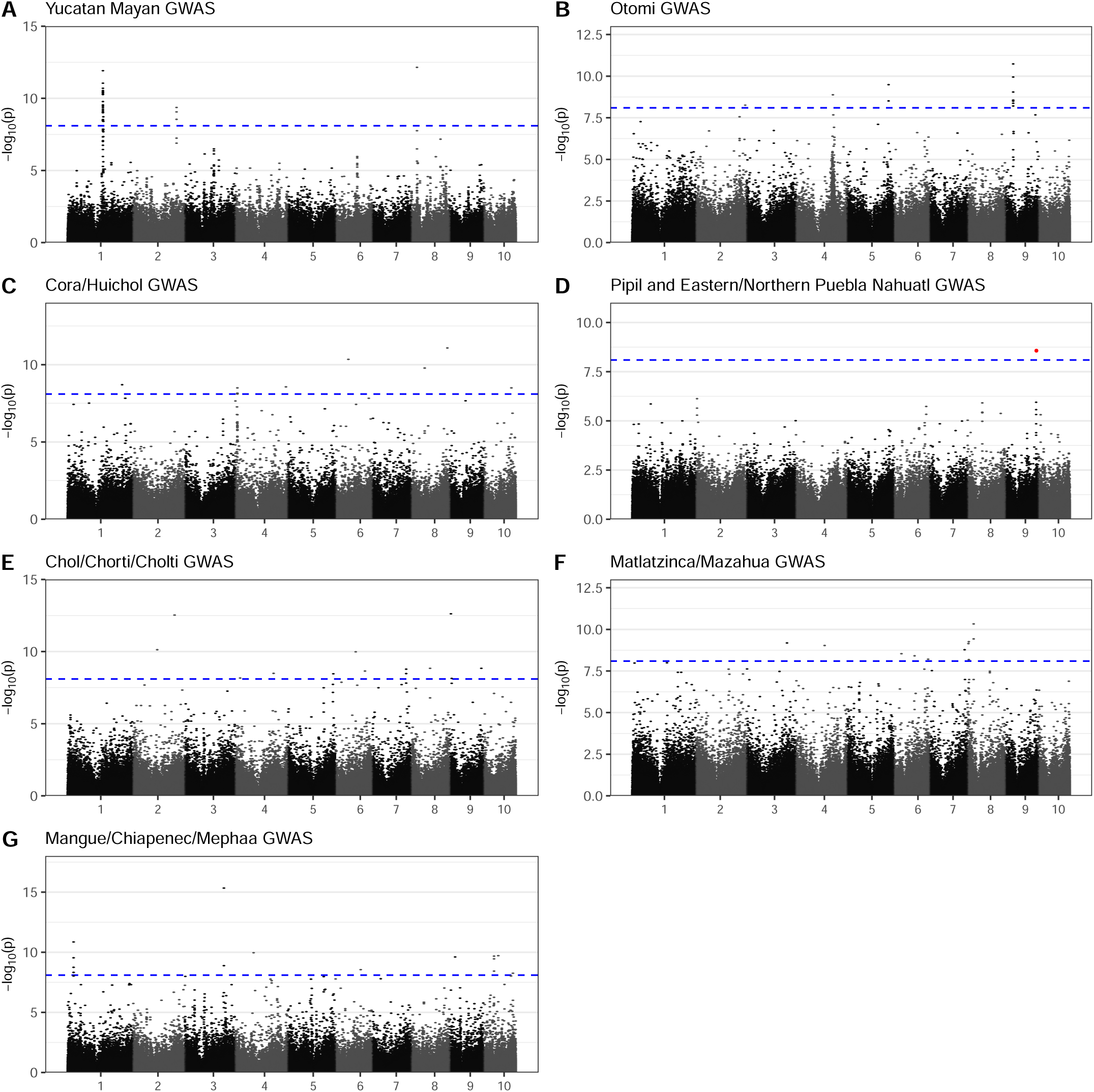
All GWAS plots. GWAS plots for association to **A)** Yucatec Mayan including Yucatec Maya, Mopan Maya, and Itza’; **B)** Querétaro Otomí, Tilapa Otomí, Mezquital Otomí, and Ixtenco Otomí; **C)** Cora and Huichol; **D)** Pipil, Eastern Nahautl, and Northern Puebla Nahuatl; **E)** Ch’ol, Chontal, Ch’orti’, and Ch’olti’; **F** San Francisco Matlatzinca, Atzingo Matlatzinca, and Michoacán Mazahua; **G)** Mangue, Chiapenec, and Mephaa. Blue dotted line indicates level of significance at 1 *×* 10*^−^*^8^.

**Figure S11.**
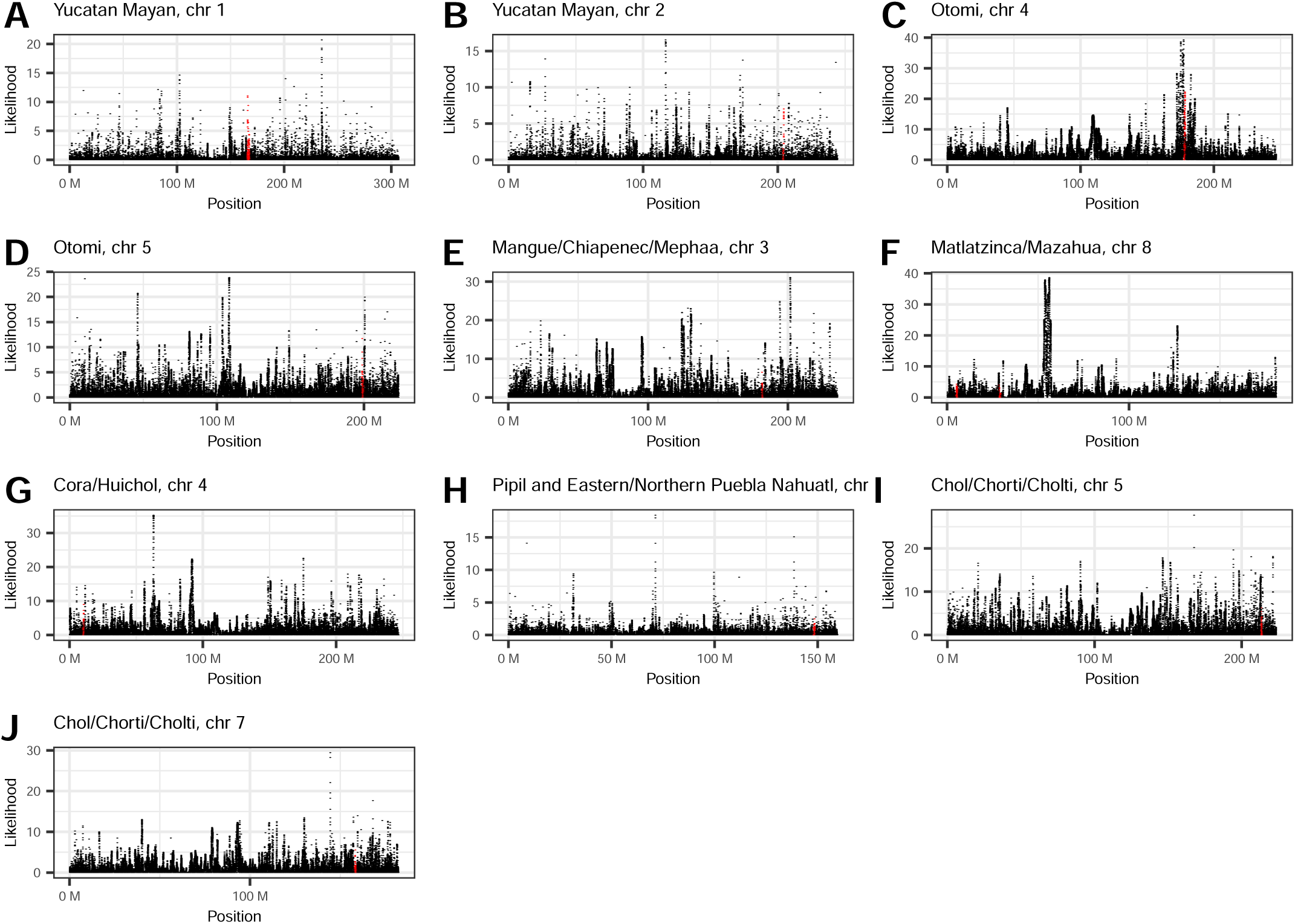
SweeD selection scan plots. SweeD plots for each linguistic sub-family in which GWAS hits were found, limited to the individual chromosome. Highlighted in red is the 1Mb region flanking the position of the top SNP in the GWAS analysis. See Fig S10 for full descriptions of linguistic sub-groupings in each panel.

**Figure S12.**
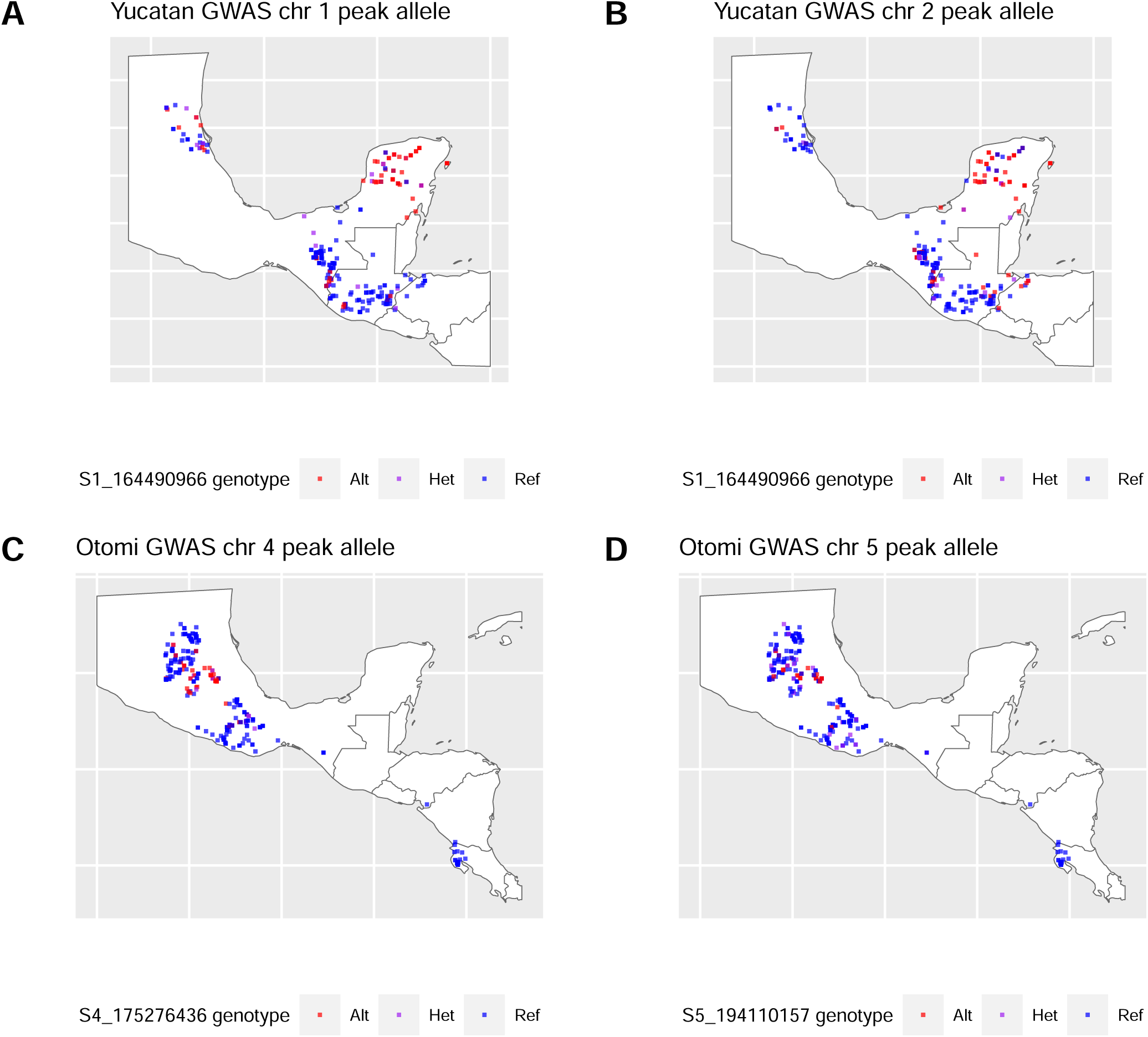
Distribution of alleles correlated with ethnolinguistics. Distribution of top GWAS peak alleles for the **A)** Yucatec Mayan, **B)** Otomí, and **C)** Cora/Huichol sub-families.

