## Supplementary figures and images for "Maize genetic diversity is largely unstructured by human ethnolinguistic diversity in its center of origin"

### Figure S8

Mayan Haynie

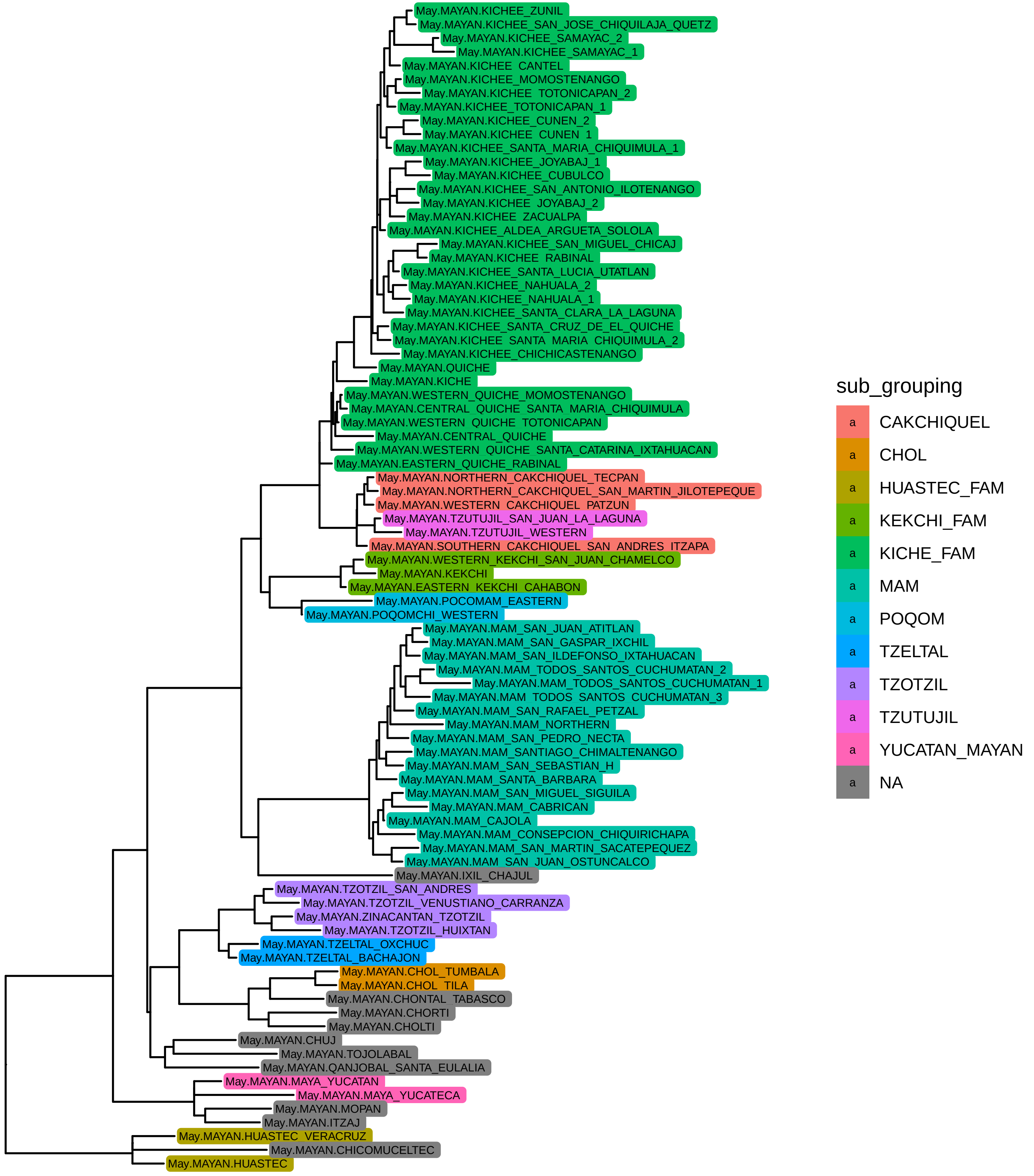

Aztecan Haynie

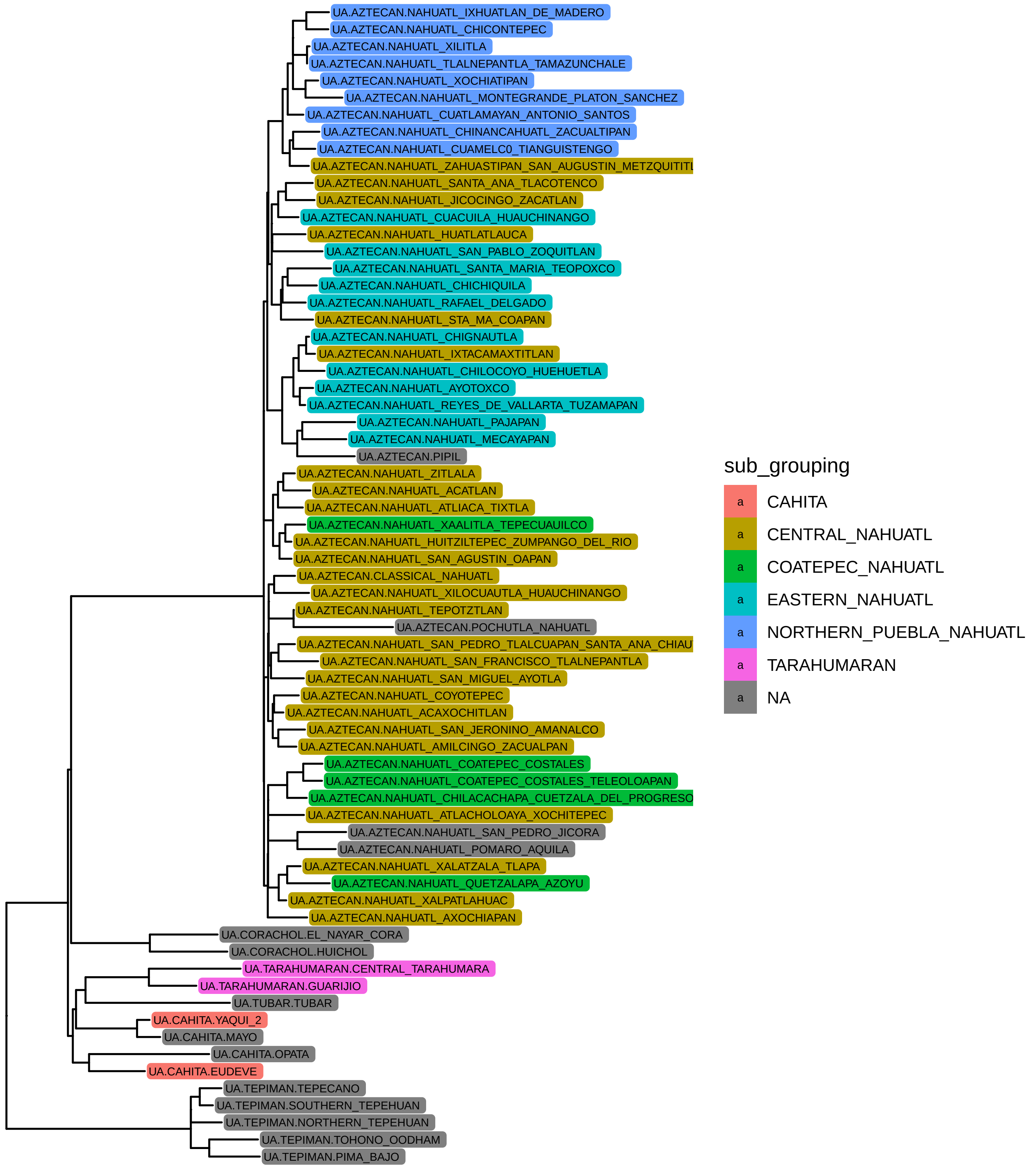

Otomanguean Haynie

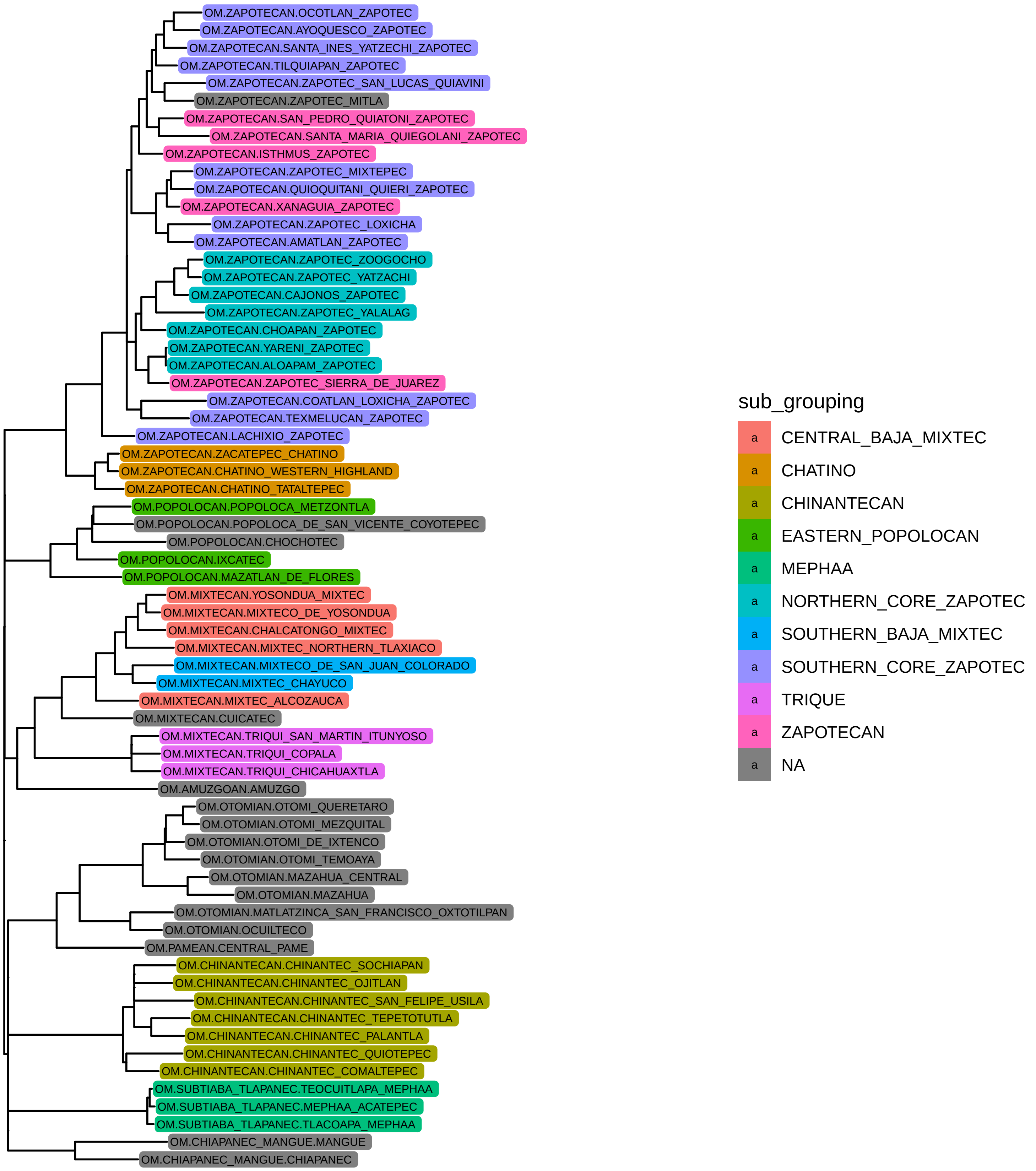

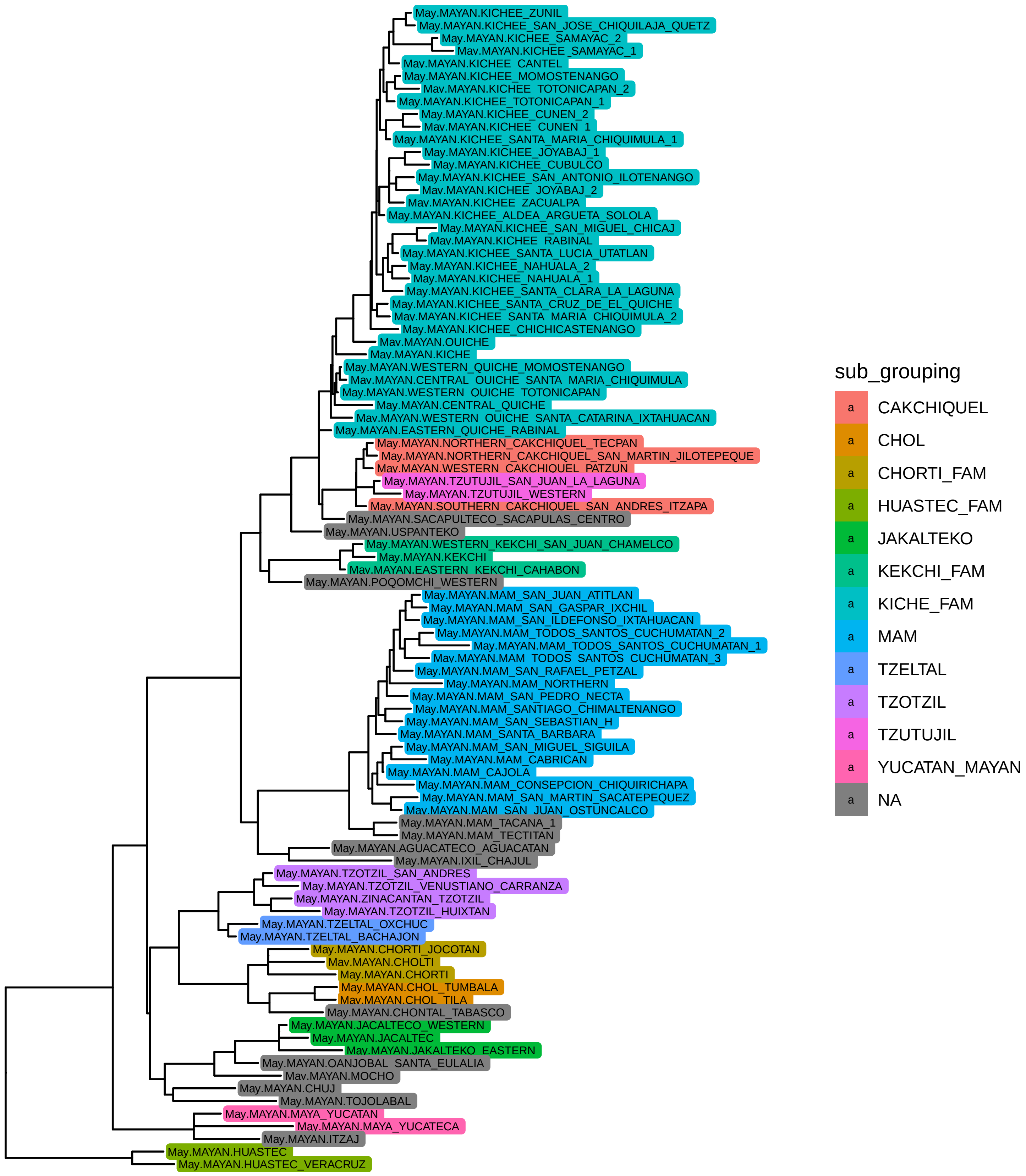

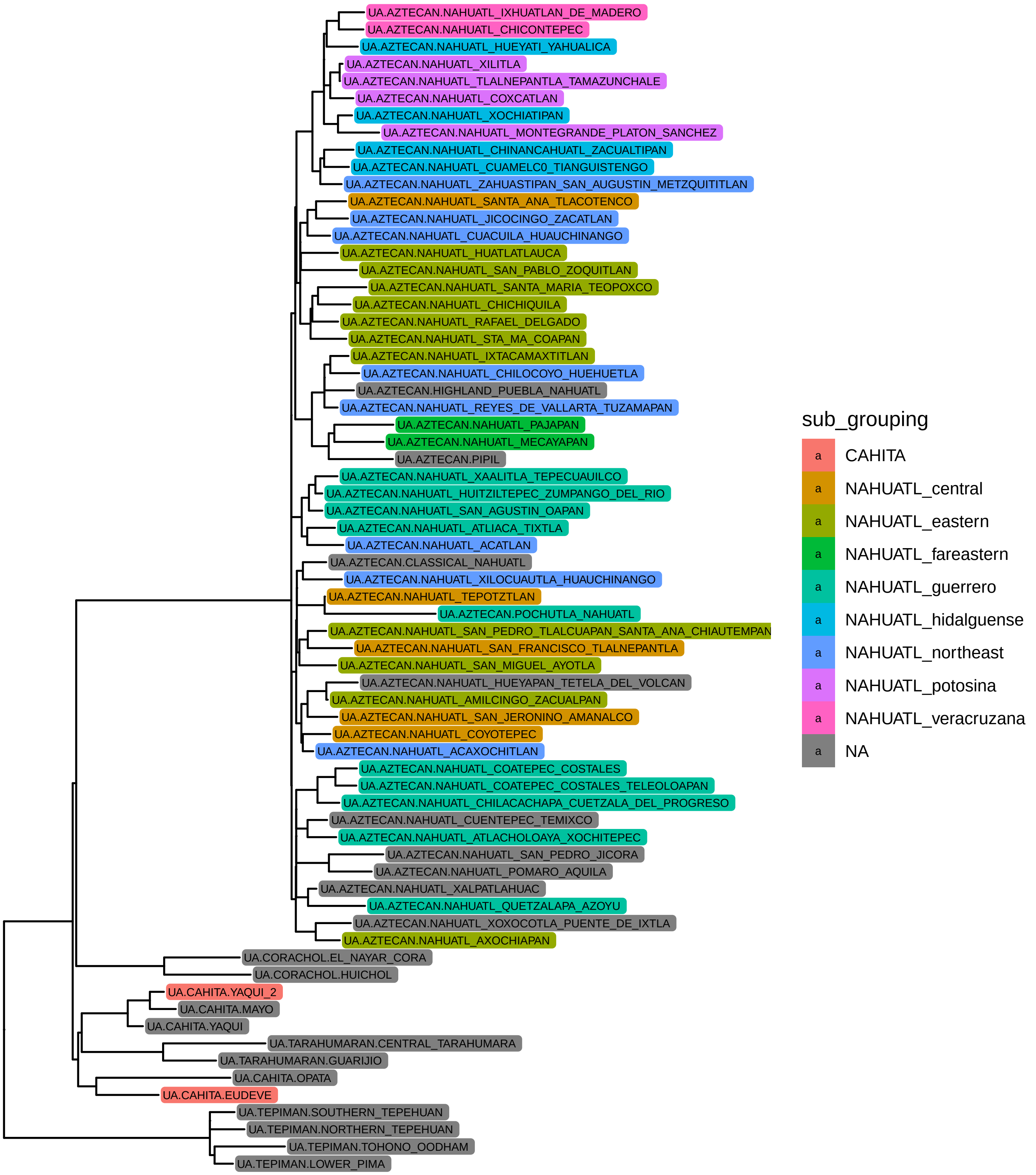

Otomanguean NL

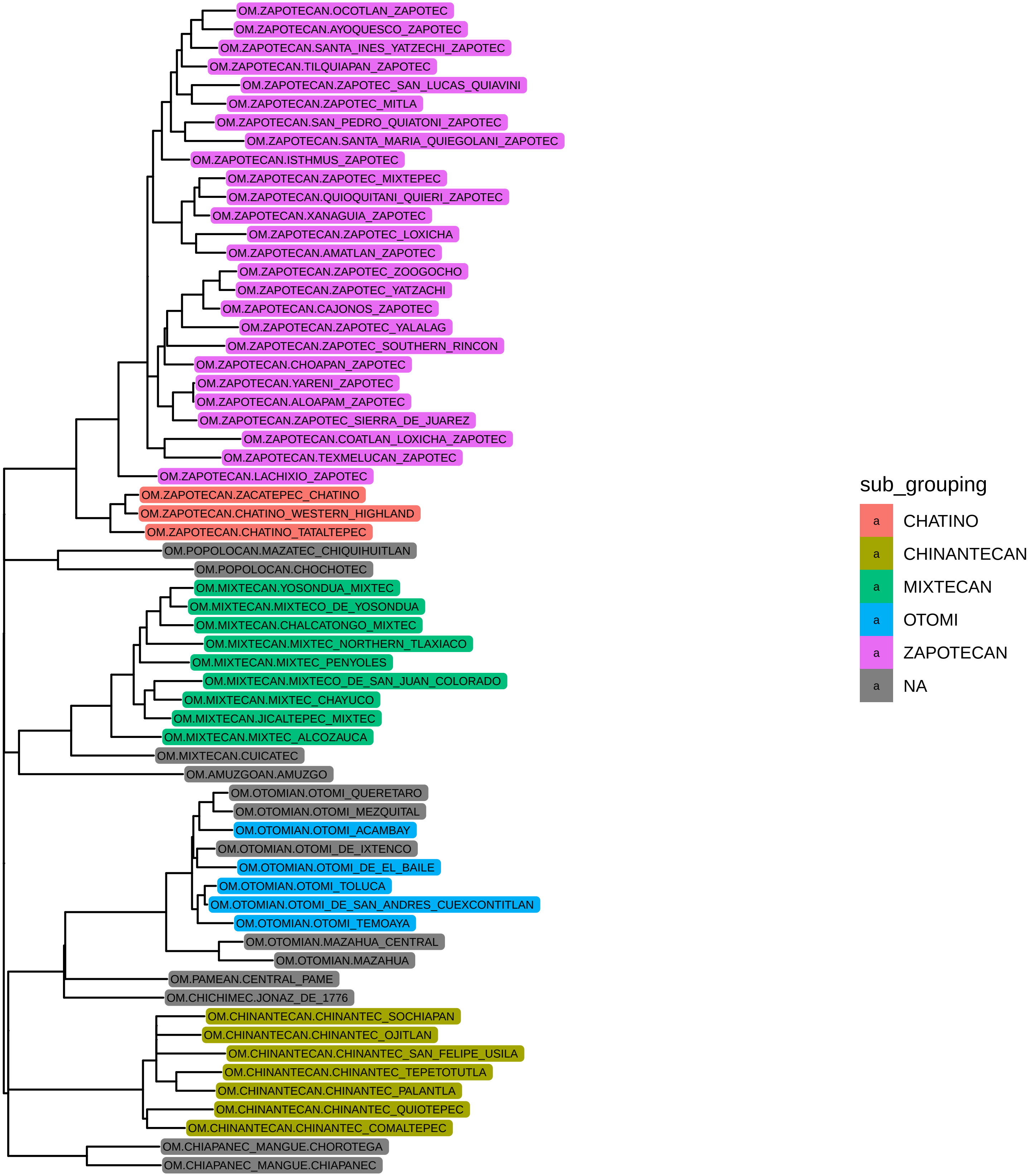

### Figure S9

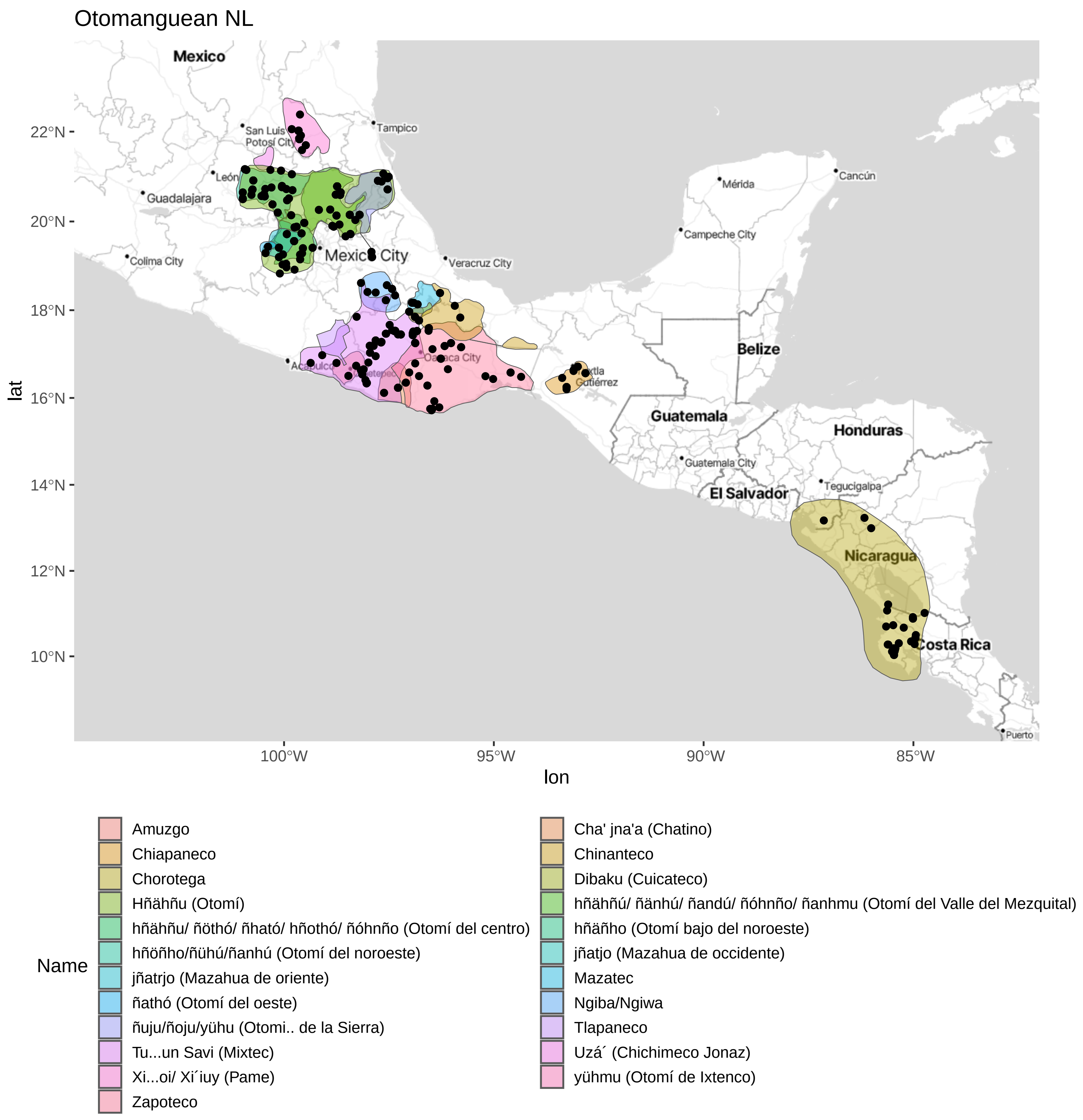

Otomanguean Haynie

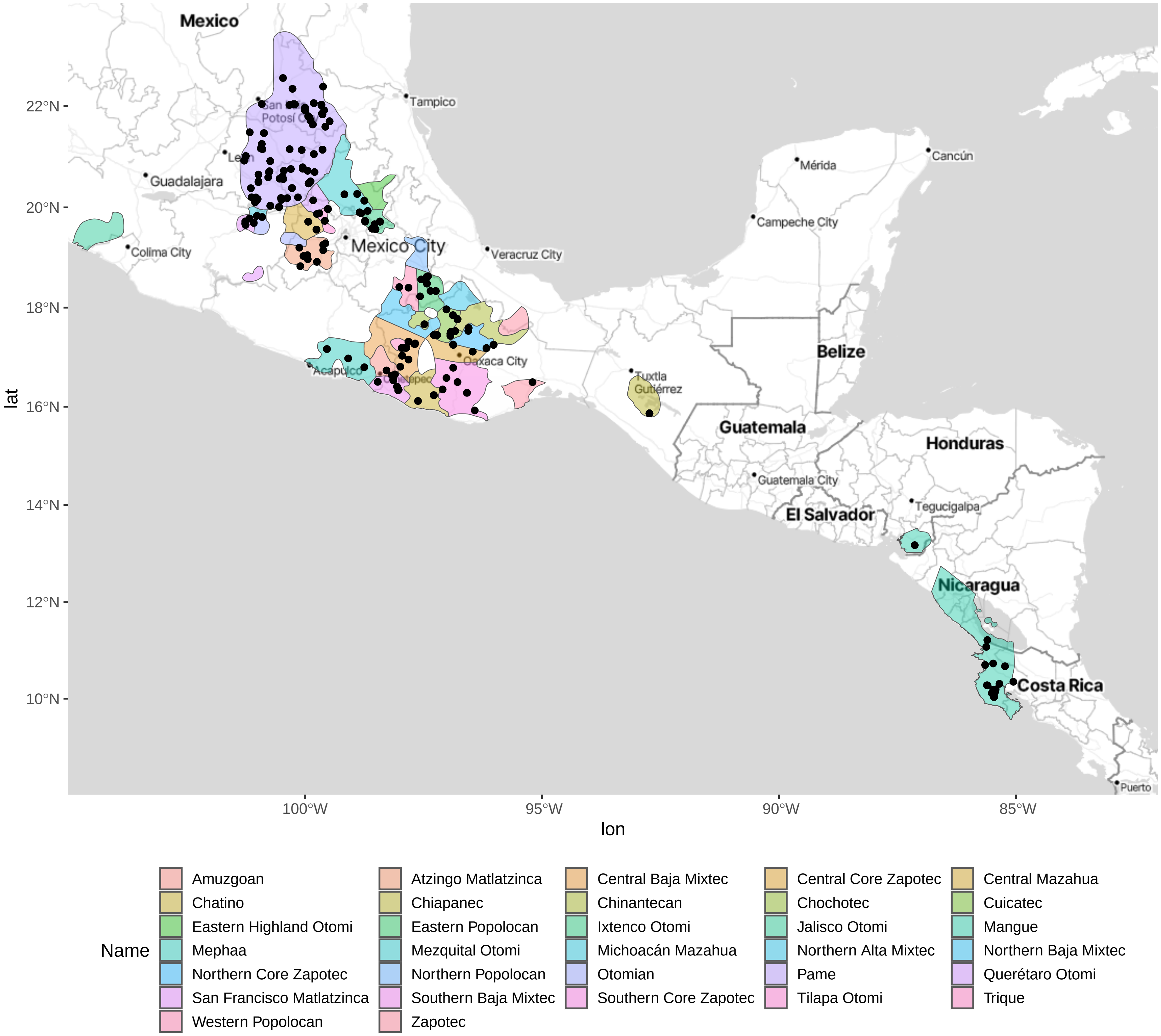

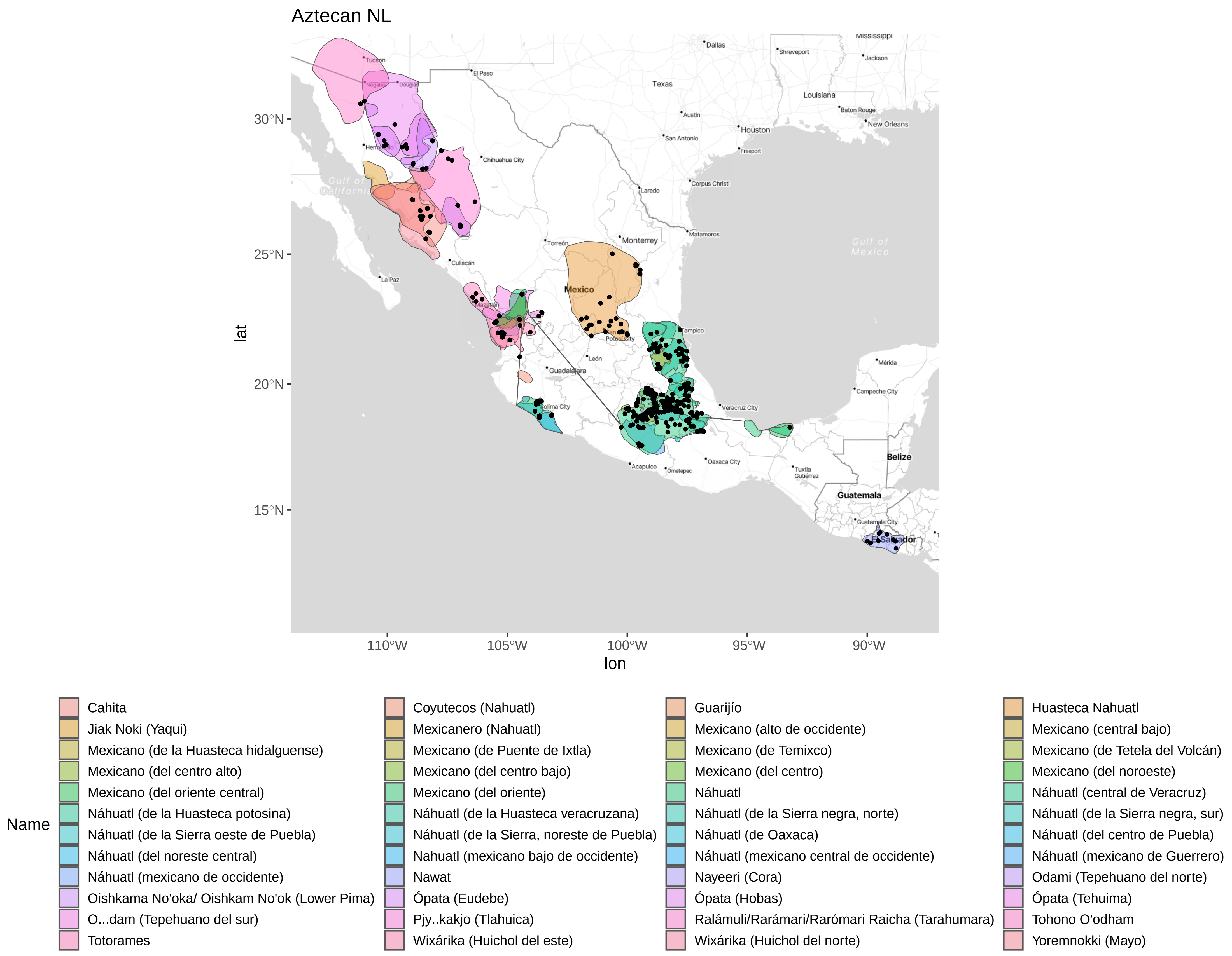

Aztecan Haynie

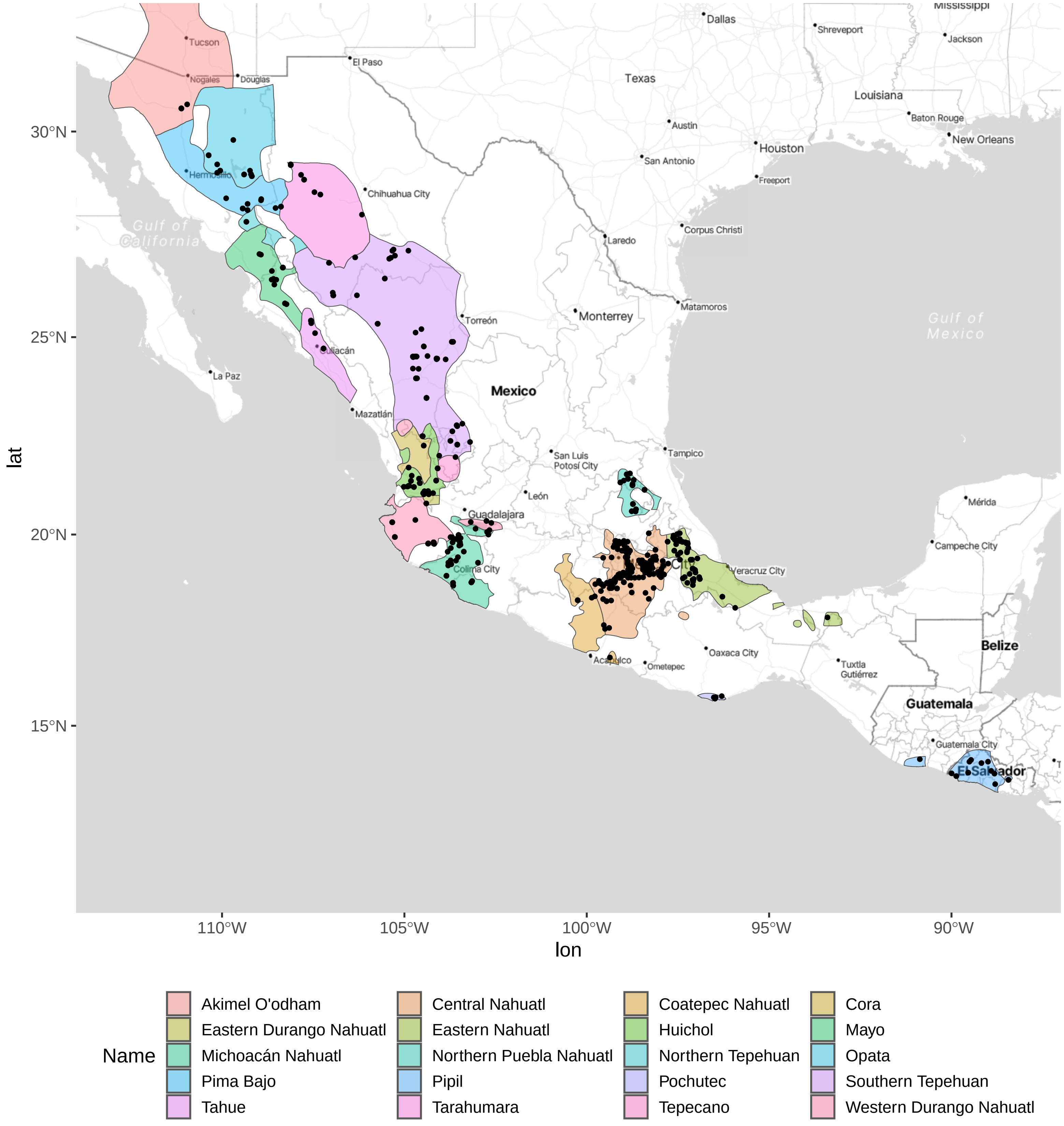

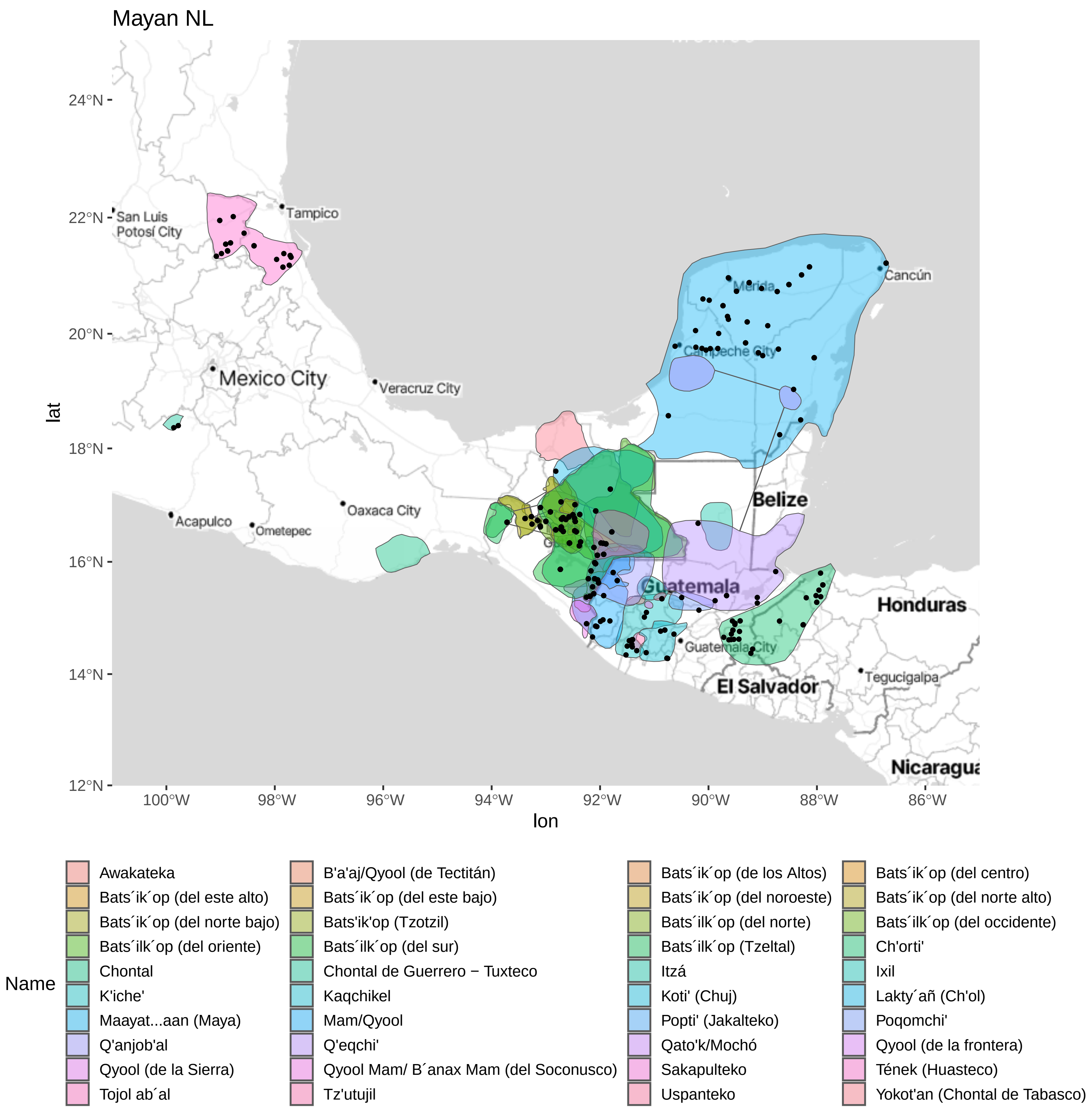

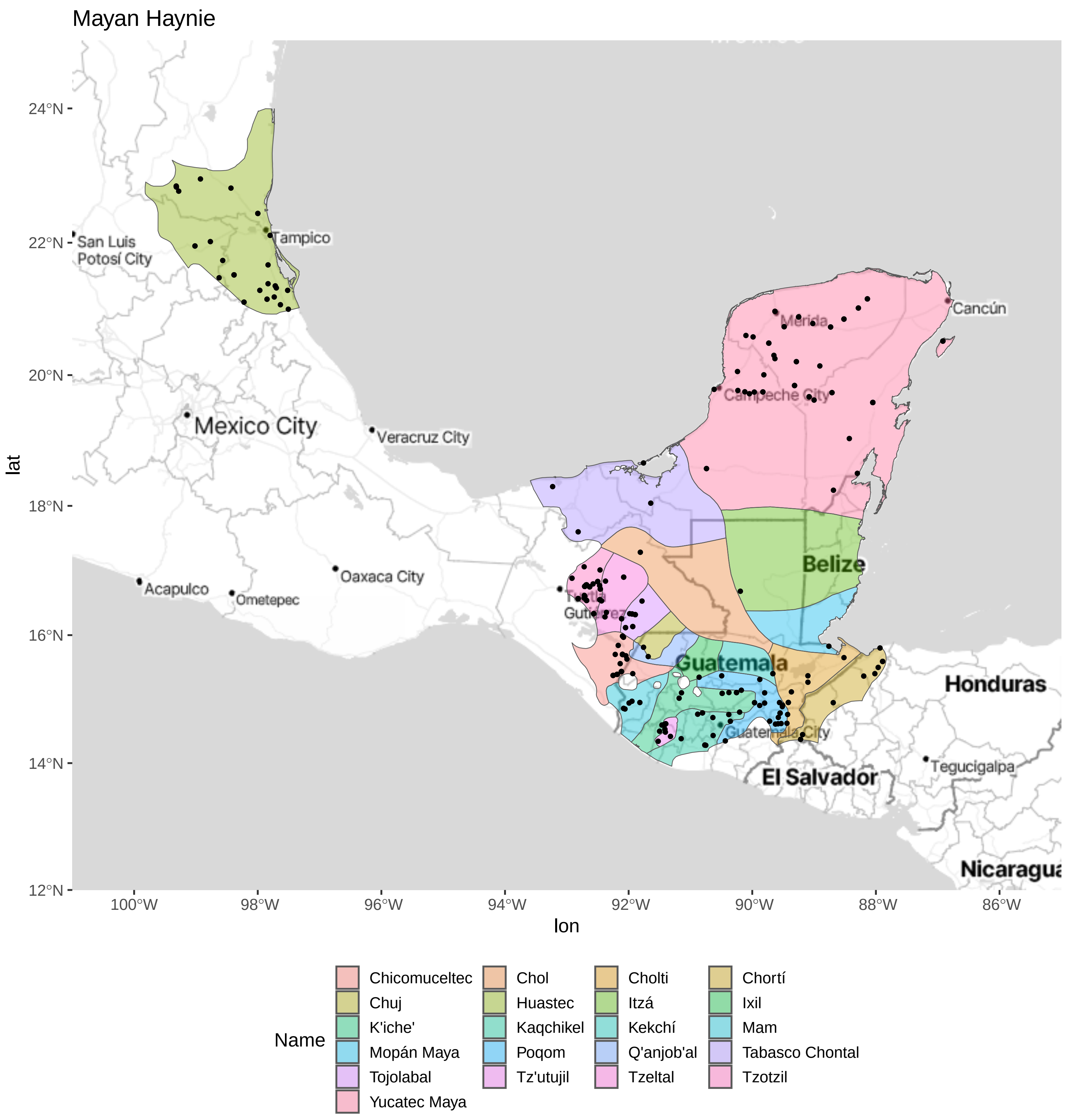
