## Supplementary material for "Maize genetic diversity is largely unstructured by human ethnolinguistic diversity in its center of origin": Figure S3

### Map of Human Samples
Color Guide

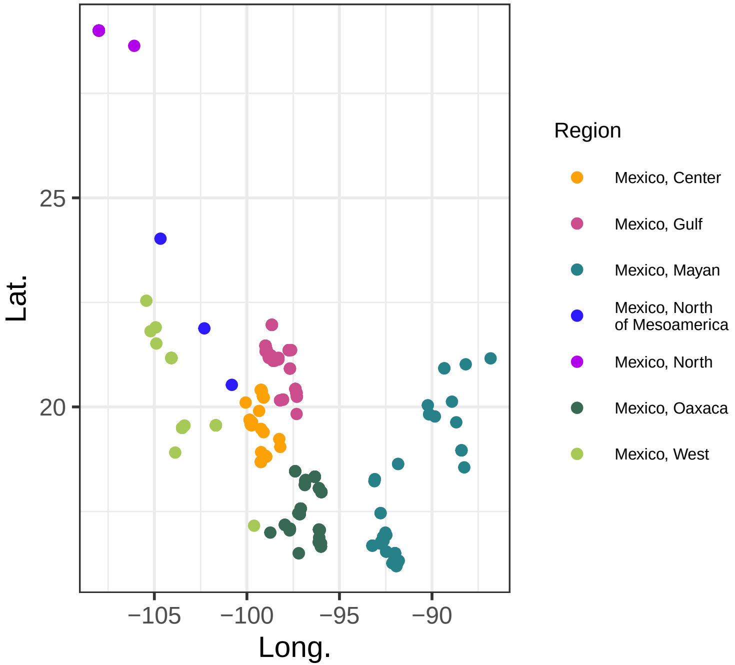

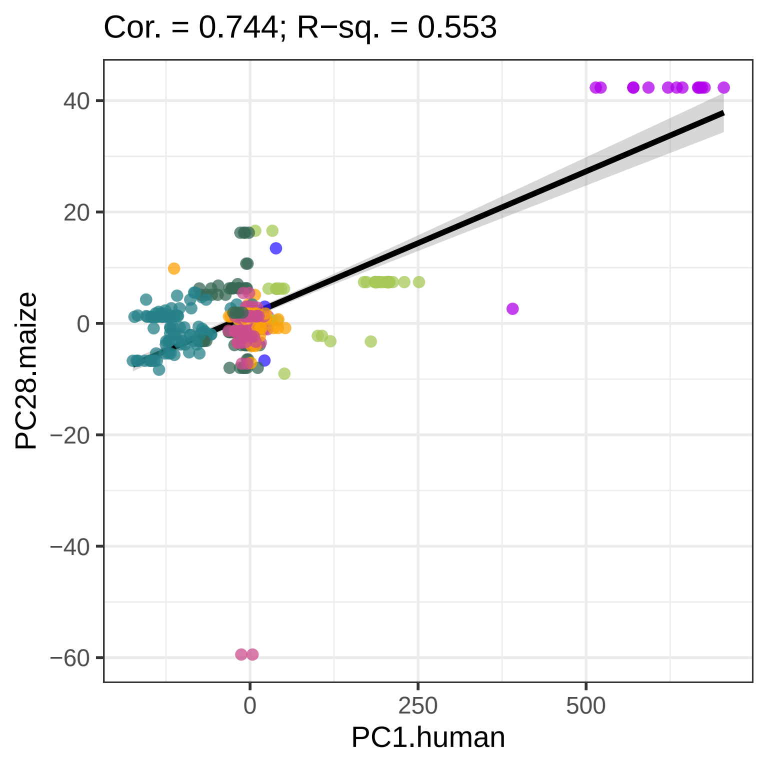
Cor. = 0.744; R−sq. = 0.553
40
20
0
−20
−40
−60
PC28.maize
0
250
500
PC1.human

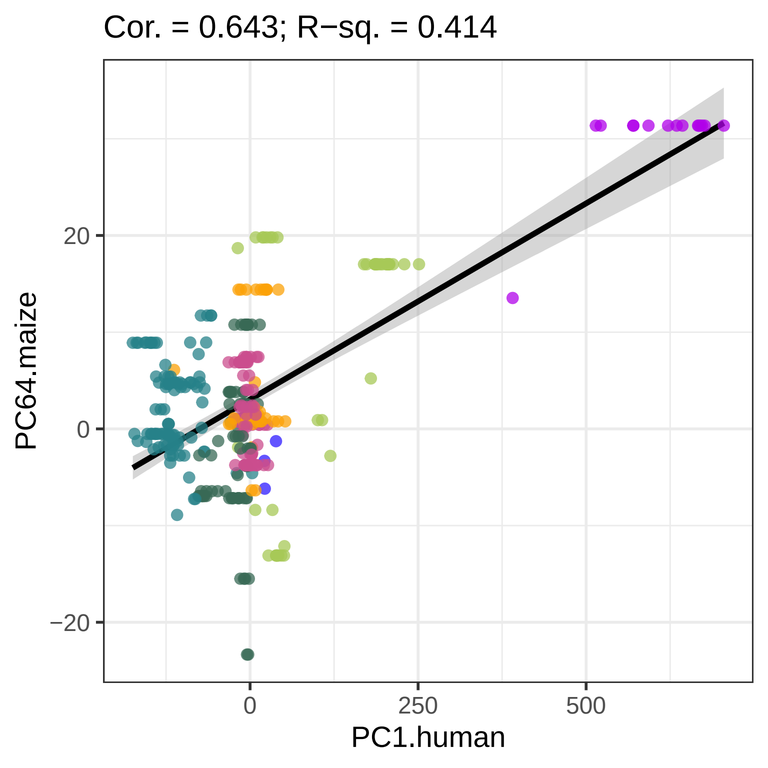

Cor. = 0.643; R−sq. = 0.414
PC64.maize
PC1.human
20
0
−20
0
250
500

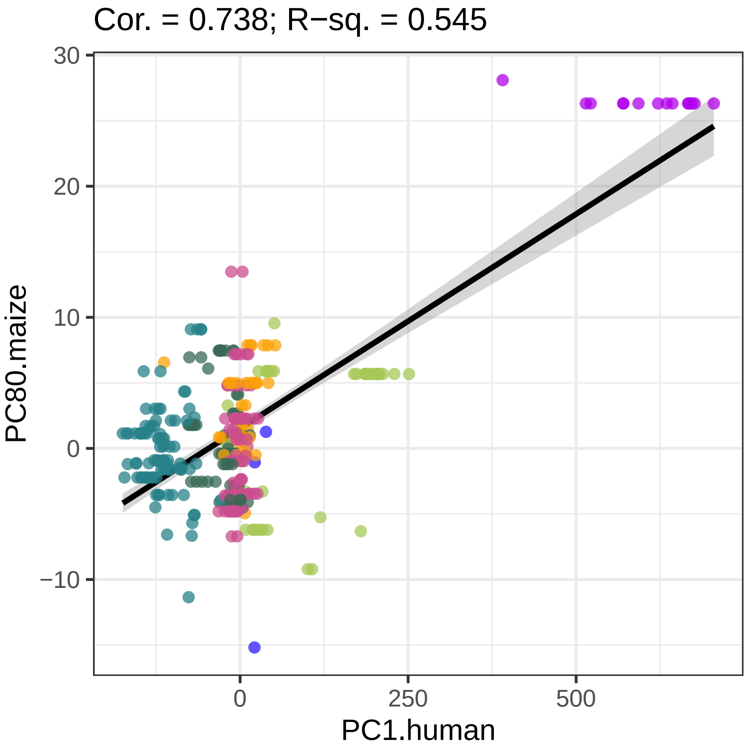

Cor. = 0.738; R−sq. = 0.545
30
20
10
0
−10
PC80.maize
0
250
500
PC1.human

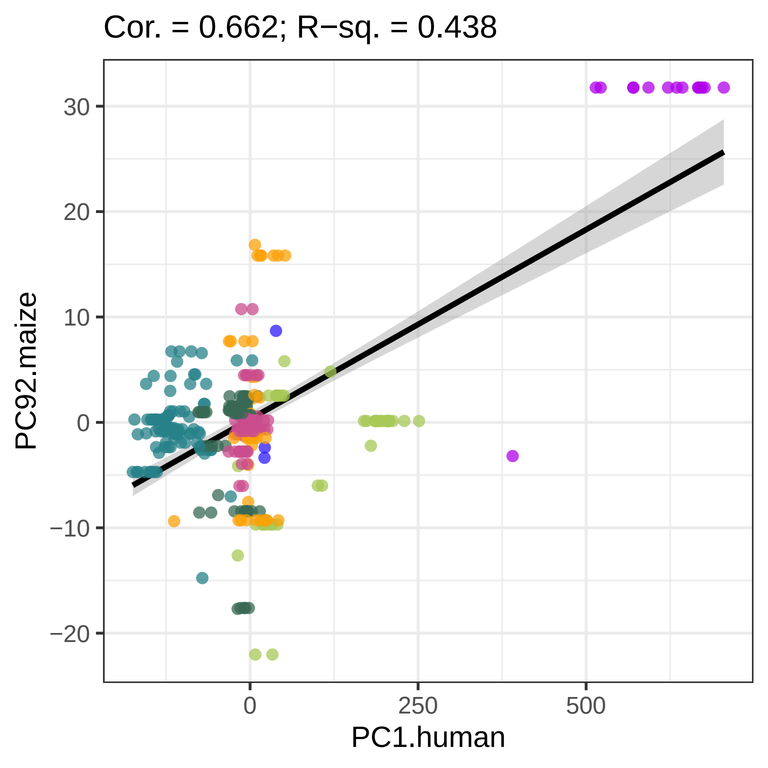
Cor. = 0.662; R−sq. = 0.438
30
20
10
0
−10
−20
PC92.maize
0
250
500
PC1.human

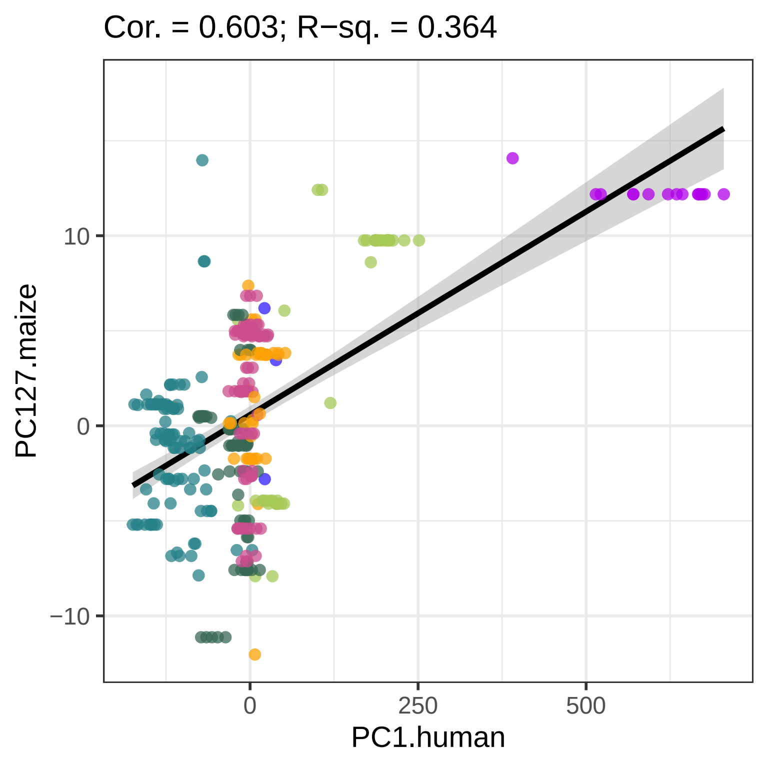
Cor. = 0.603; R−sq. = 0.364
PC127.maize
PC1.human
10
0
−10
0
250
500

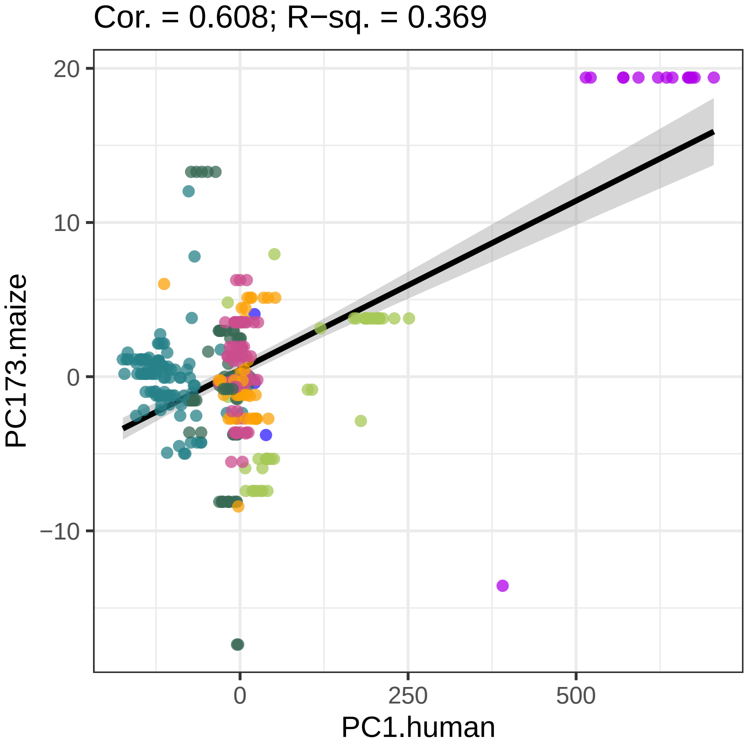

Cor. = 0.608; R−sq. = 0.369
PC173.maize
PC1.human
20
10
0
−10
0
250
500

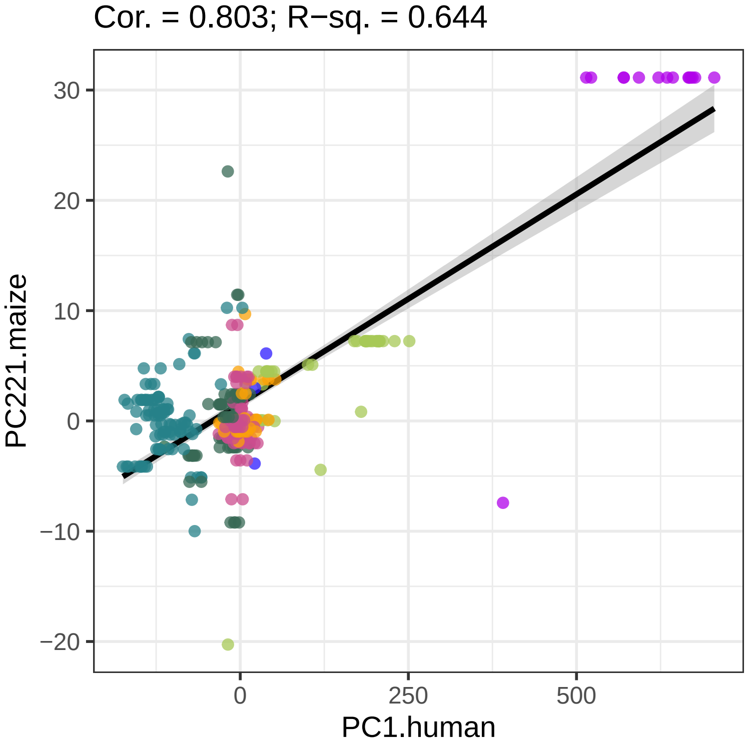

Cor. = 0.803; R−sq. = 0.644
PC221.maize
PC1.human
30
20
10
0
−10
−20
0
250
500

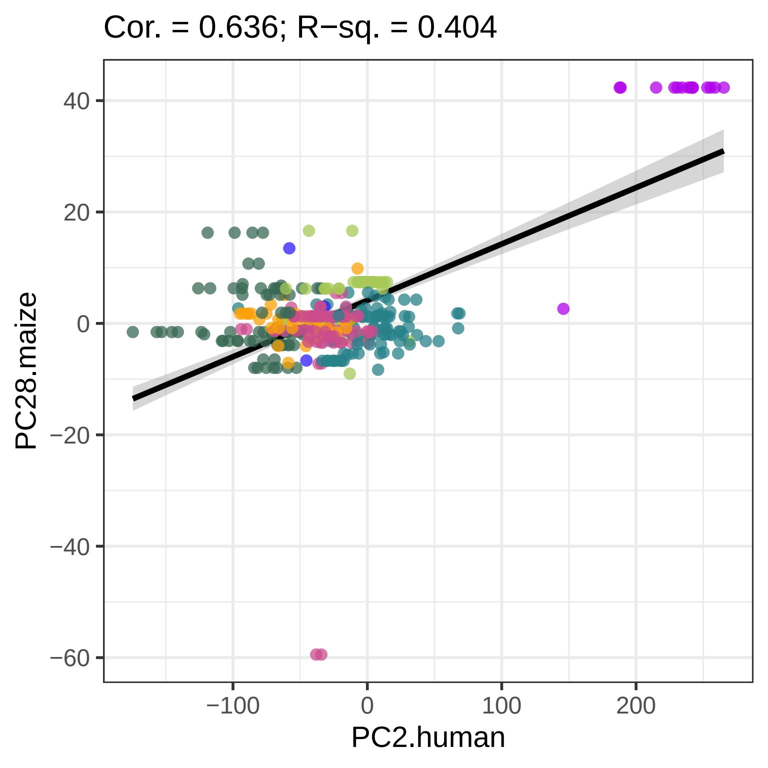
Cor. = 0.636; R−sq. = 0.404
PC28.maize
PC2.human
40
20
0
−20
−40
−60
−100
0
100
200

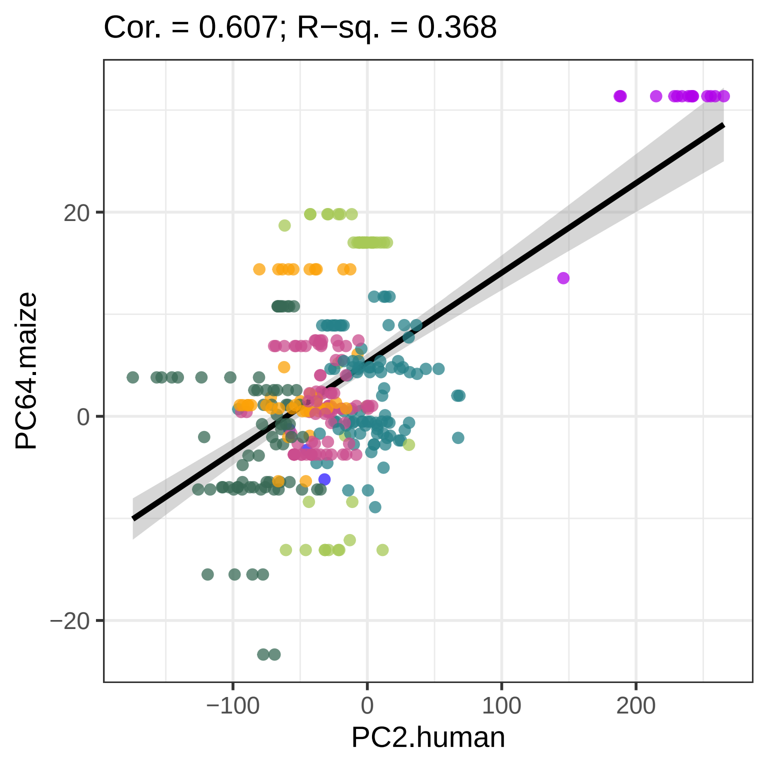
Cor. = 0.607; R−sq. = 0.368
PC64.maize
PC2.human
20
0
−20
−100
0
100
200

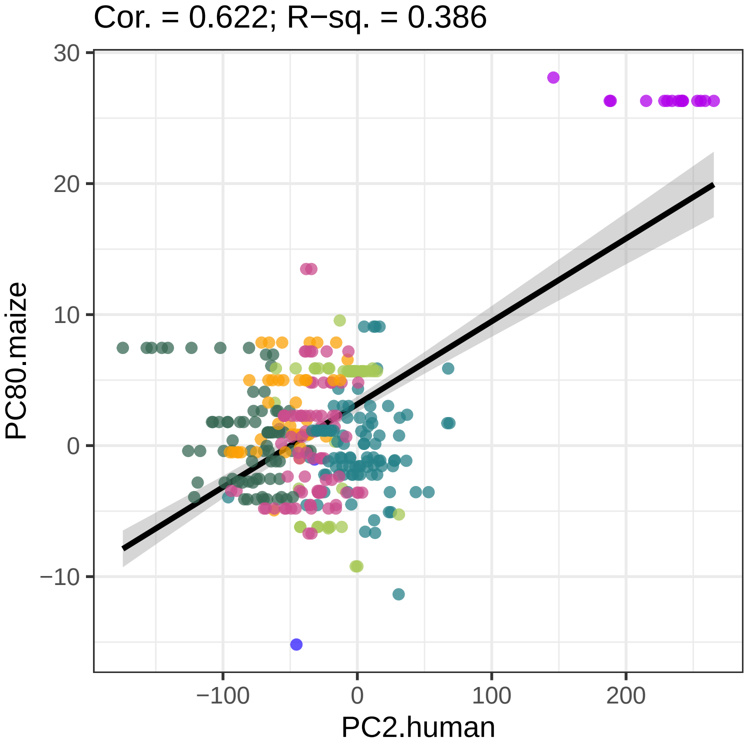
Cor. = 0.622; R−sq. = 0.386
30
20
10
0
−10
PC80.maize
−100
0
100
200
PC2.human

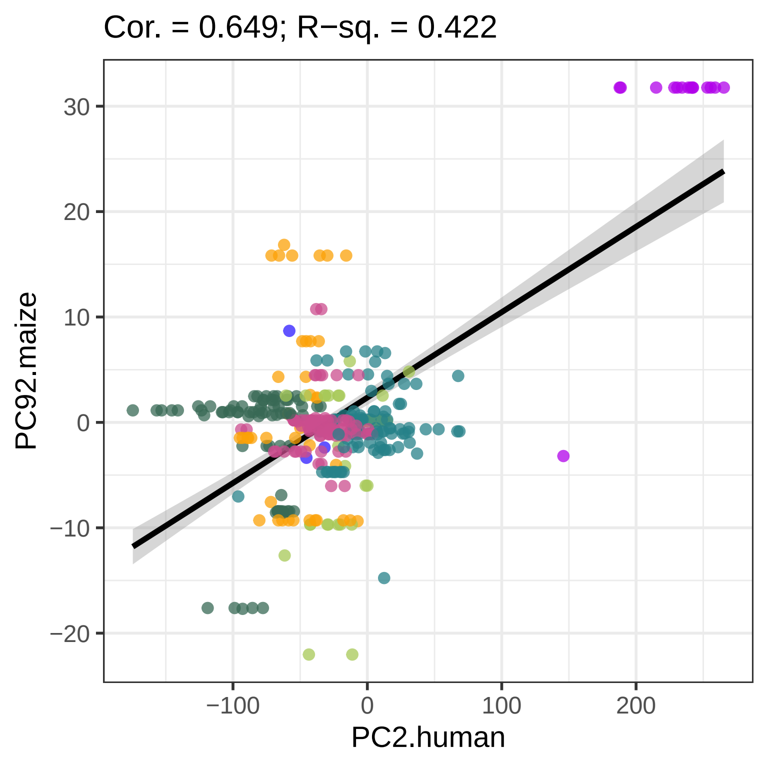

Cor. = 0.649; R−sq. = 0.422
PC92.maize
30
20
10
0
−10
−20
−100
0
100
200
PC2.human

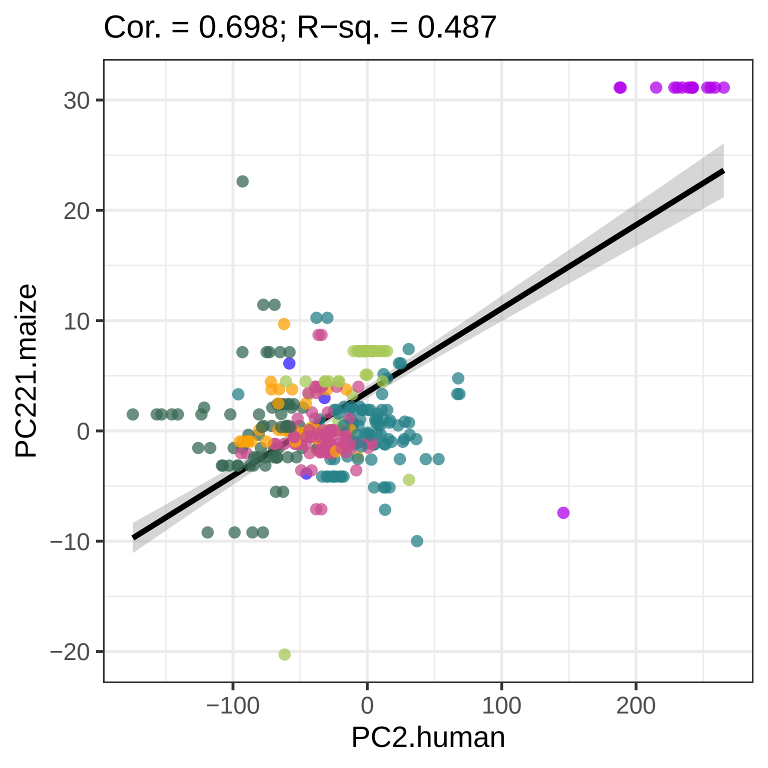

Cor. = 0.698; R−sq. = 0.487
30
20
10
0
−10
−20
PC221.maize
−100
0
100
200
PC2.human

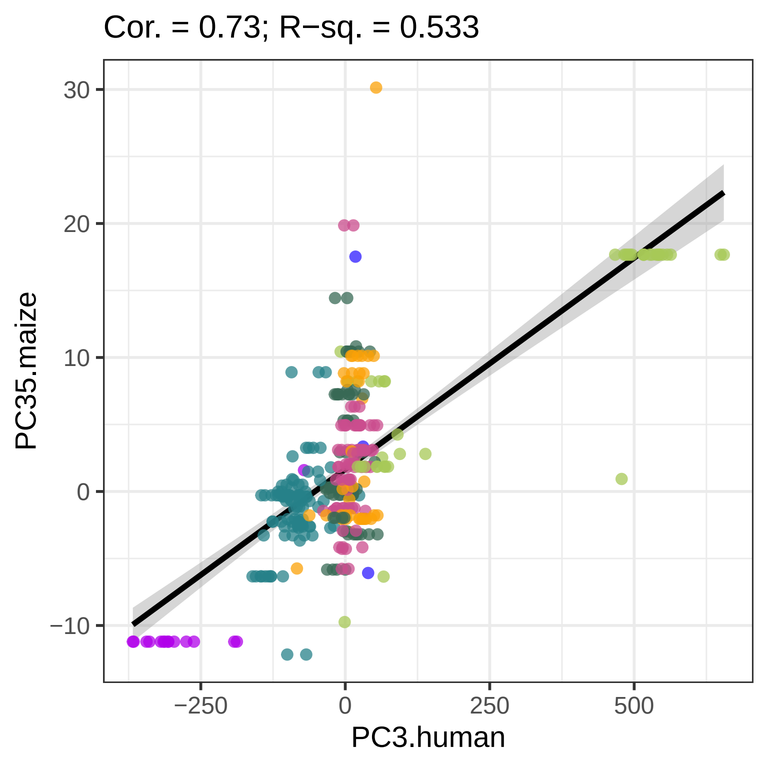
Cor. = 0.73; R−sq. = 0.533
30
20
10
0
−10
PC35.maize
−250
0
250
500
PC3.human

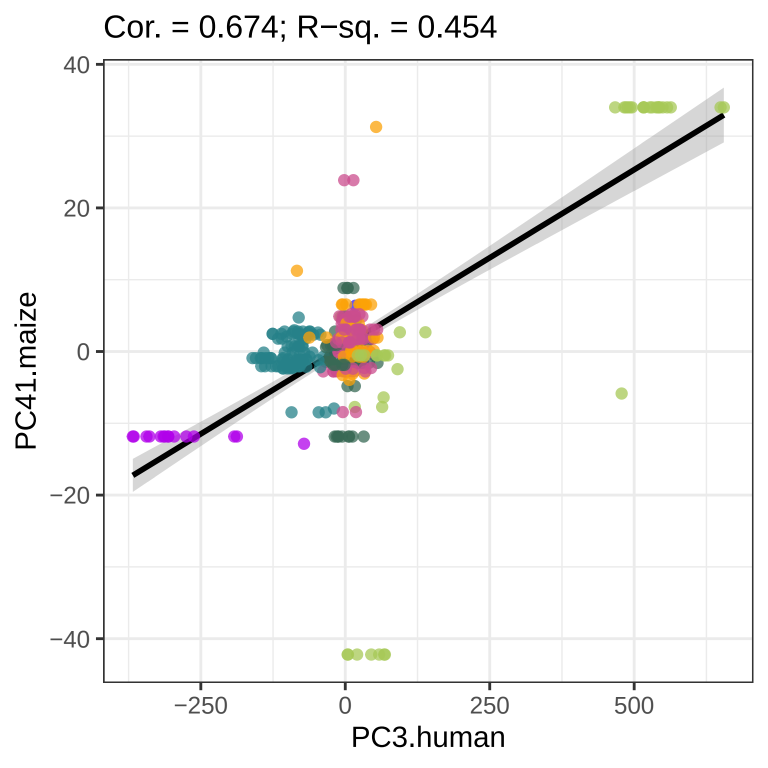
Cor. = 0.674; R−sq. = 0.454
40
20
0
−20
−40
PC41.maize
−250
0
250
500
PC3.human

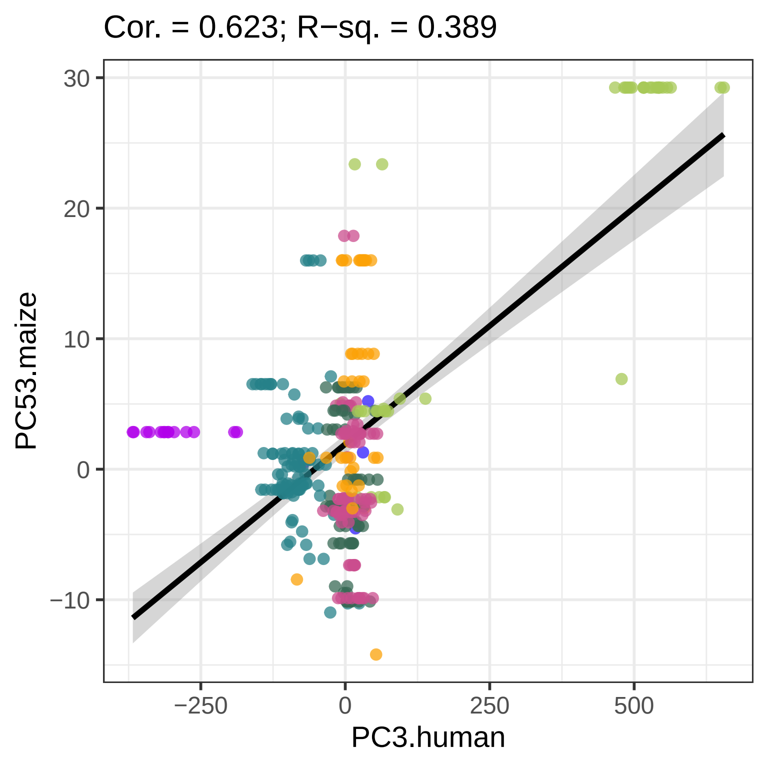
Cor. = 0.623; R−sq. = 0.389
30
20
10
0
−10
PC53.maize
−250
0
250
500
PC3.human

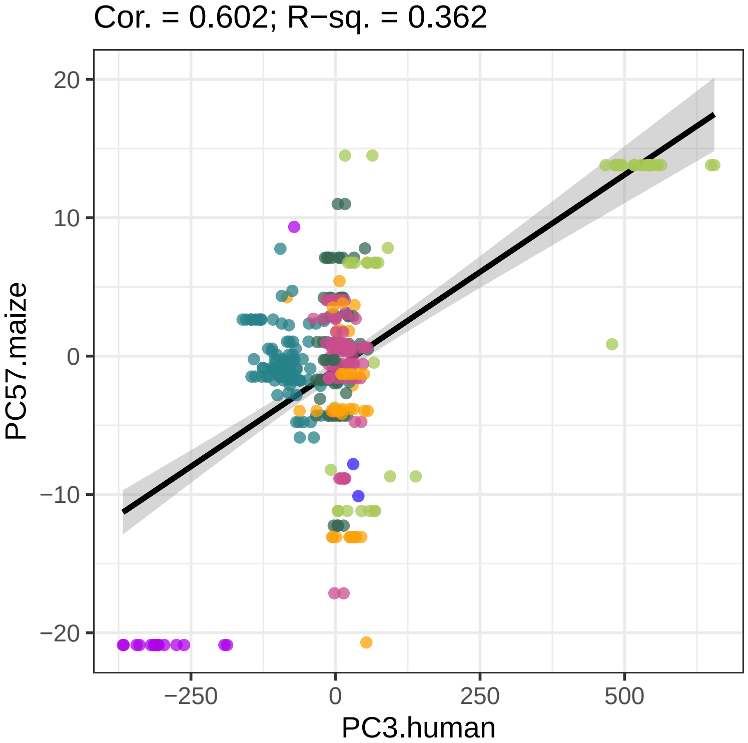
Cor. = 0.602; R−sq. = 0.362
20
10
0
−10
−20
PC57.maize
−250
0
250
500
PC3.human

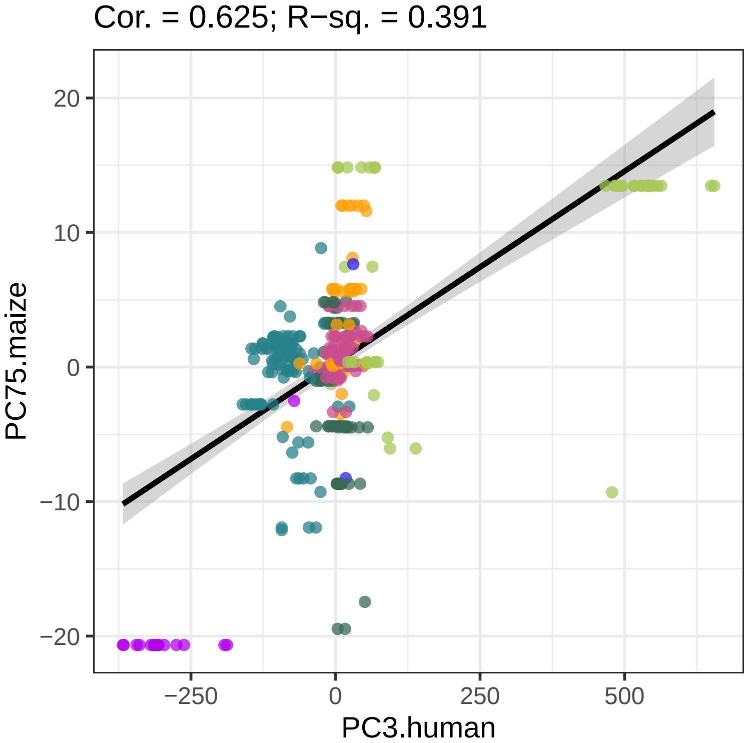
Cor. = 0.625; R−sq. = 0.391
20
10
0
−10
−20
PC75.maize
−250
0
250
500
PC3.human

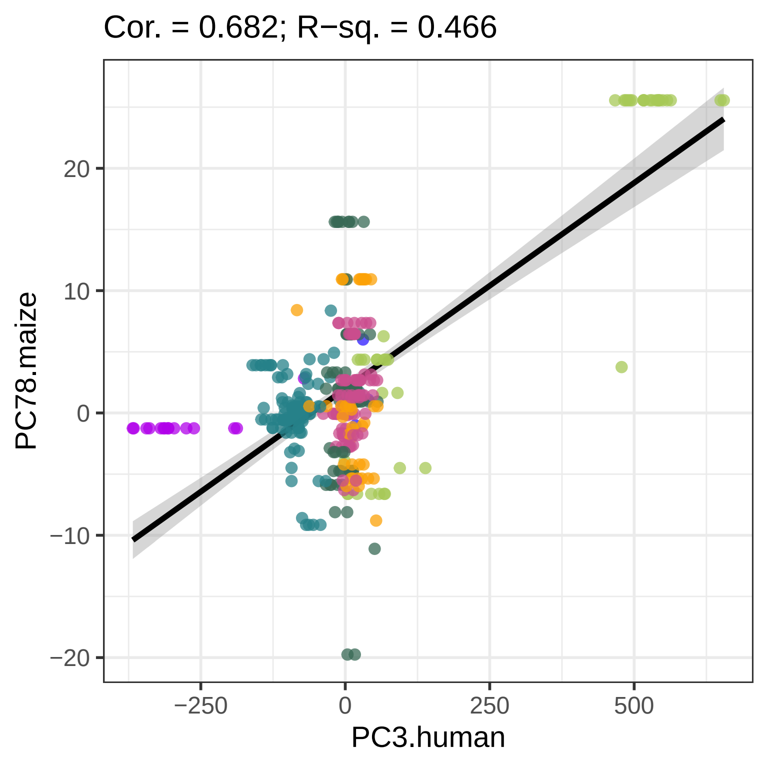
Cor. = 0.682; R−sq. = 0.466
PC78.maize
PC3.human
20
10
0
−10
−20
−250
0
250
500

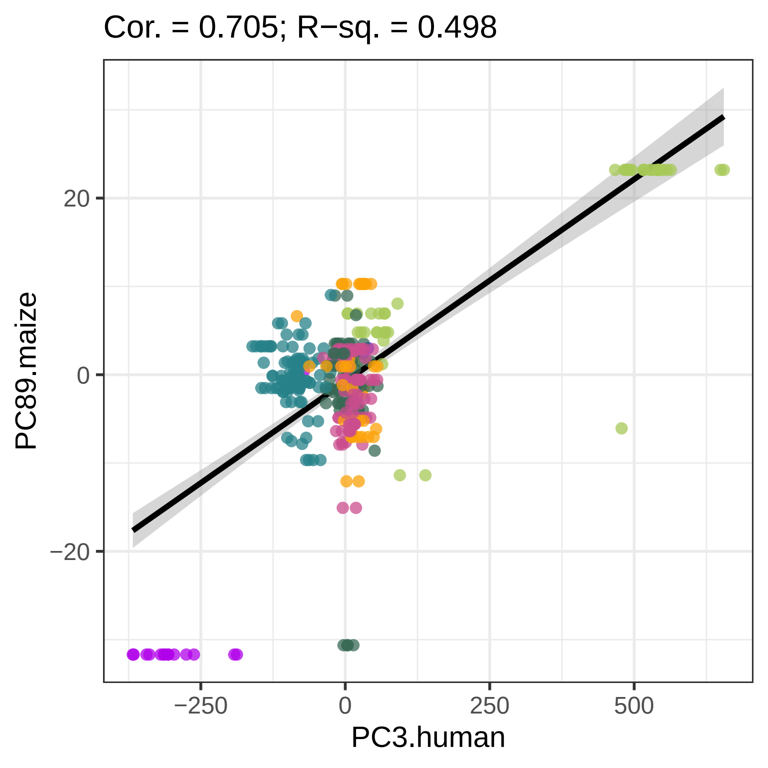
Cor. = 0.705; R−sq. = 0.498
PC89.maize
20
0
−20
−250
0
250
500
PC3.human

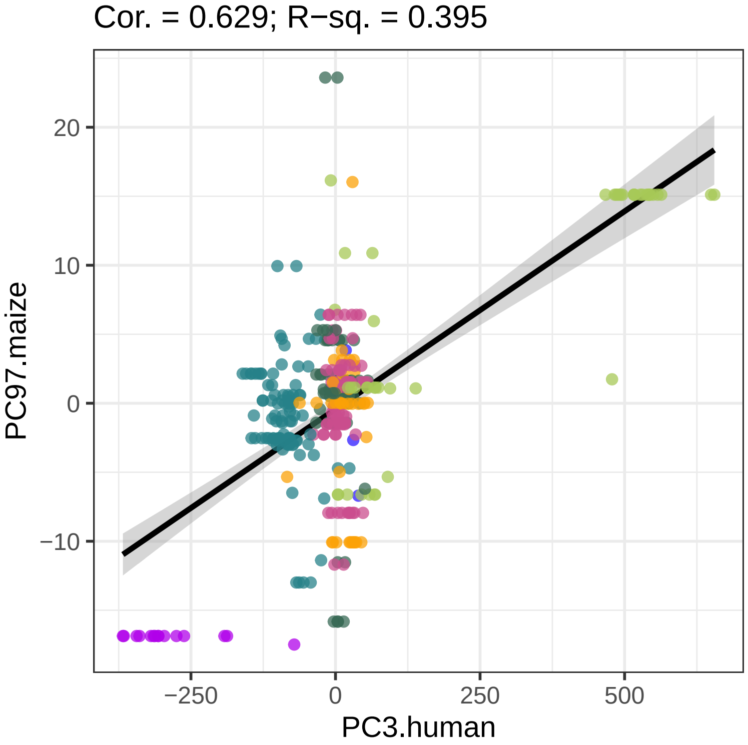
Cor. = 0.629; R−sq. = 0.395
20
10
0
−10
PC97.maize
−250
0
250
500
PC3.human

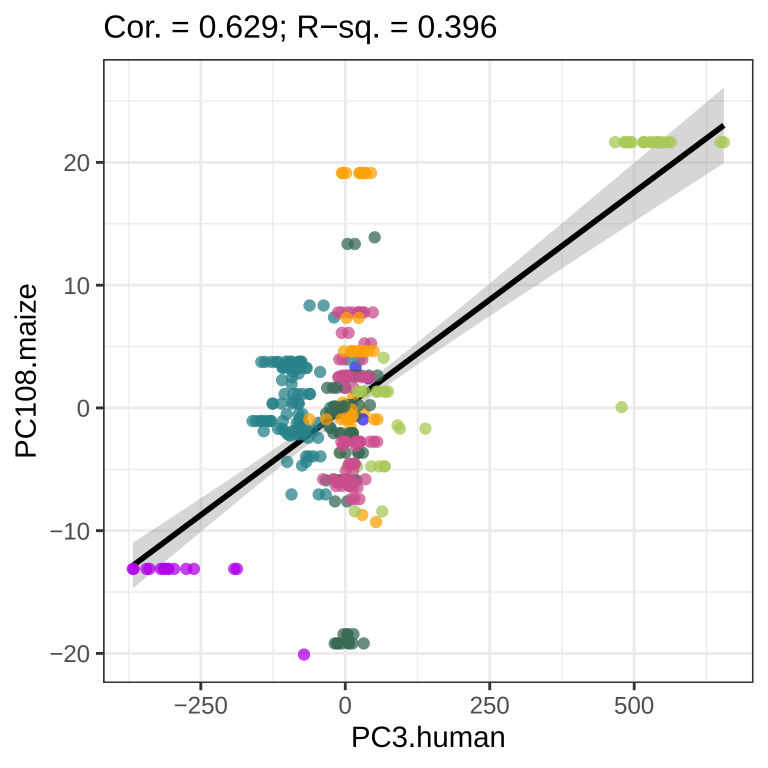

Cor. = 0.629; R−sq. = 0.396
PC108.maize
PC3.human
20
10
0
−10
−20
−250
0
250
500

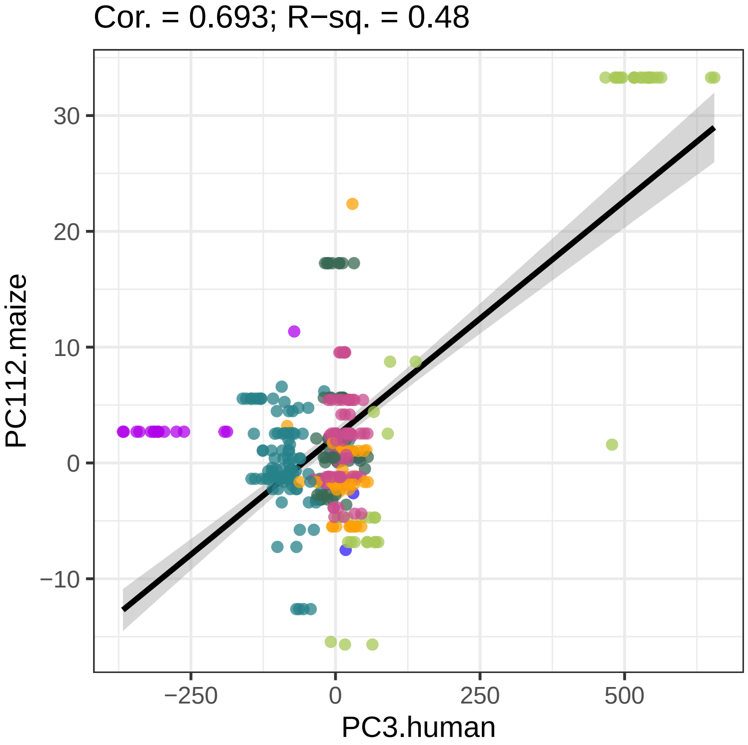
Cor. = 0.693; R−sq. = 0.48
PC112.maize
30
20
10
0
−10
−250
0
250
500
PC3.human

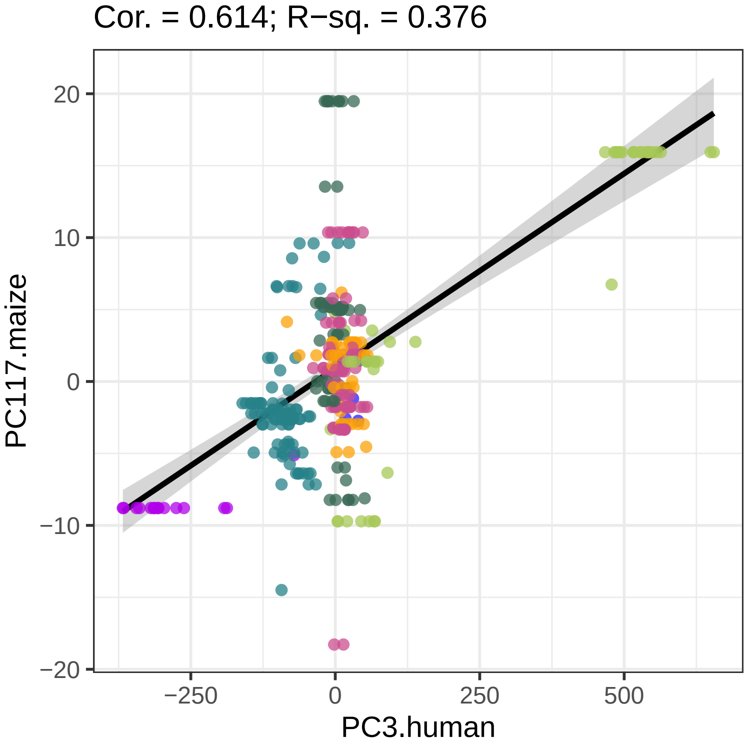
Cor. = 0.614; R−sq. = 0.376
20
10
0
−10
−20
PC117.maize
−250
0
250
500
PC3.human

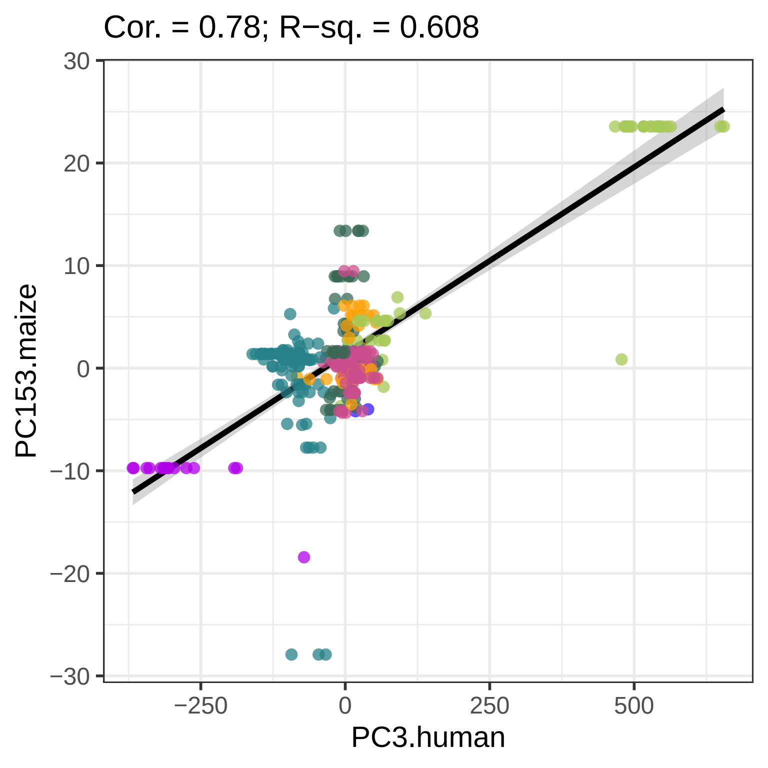
Cor. = 0.78; R−sq. = 0.608
PC153.maize
30
20
10
0
−10
−20
−30
−250
0
250
500
PC3.human

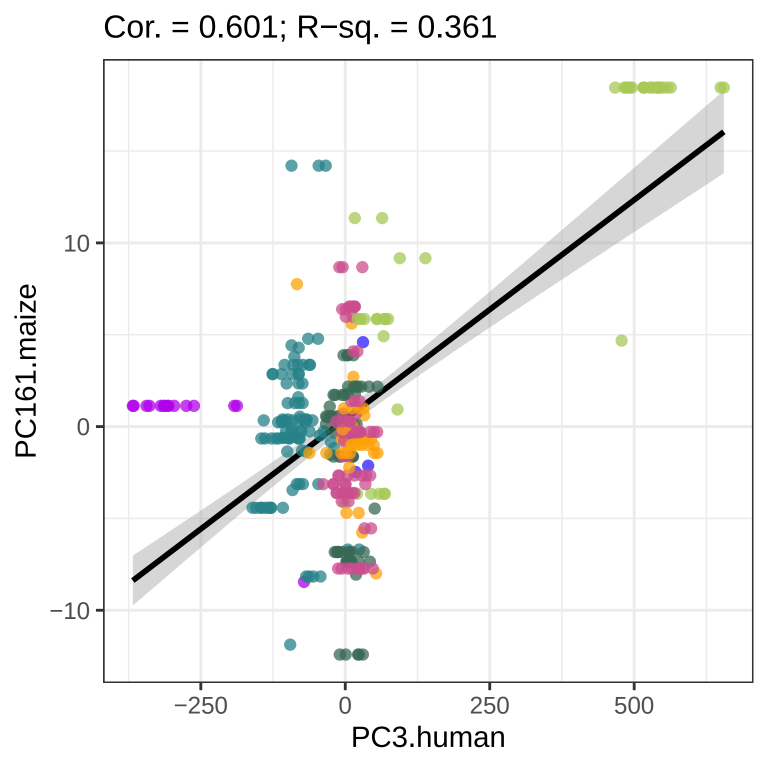

Cor. = 0.601; R−sq. = 0.361
PC161.maize
10
0
−10
−250
0
250
500
PC3.human

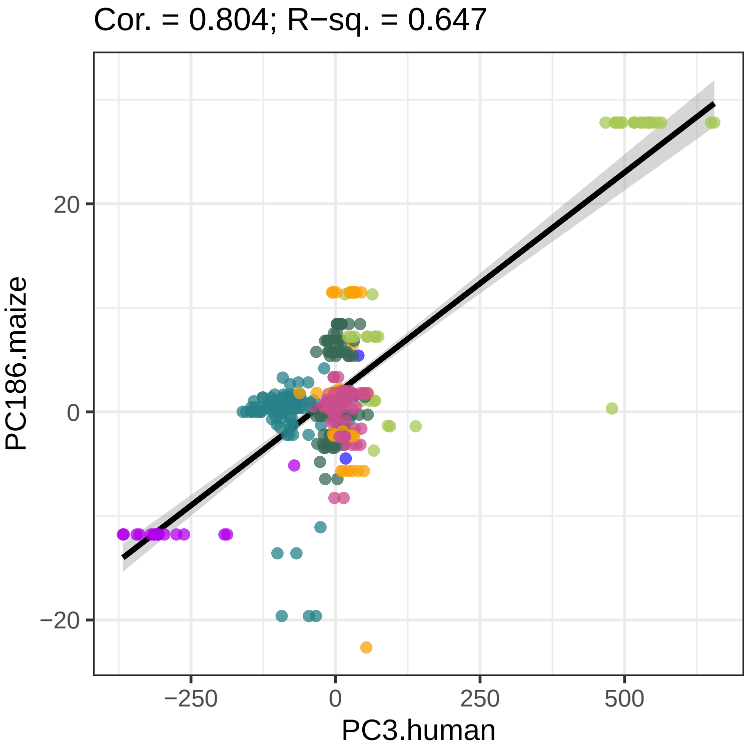

Cor. = 0.804; R−sq. = 0.647
PC186.maize
20
0
−20
−250
0
250
500
PC3.human

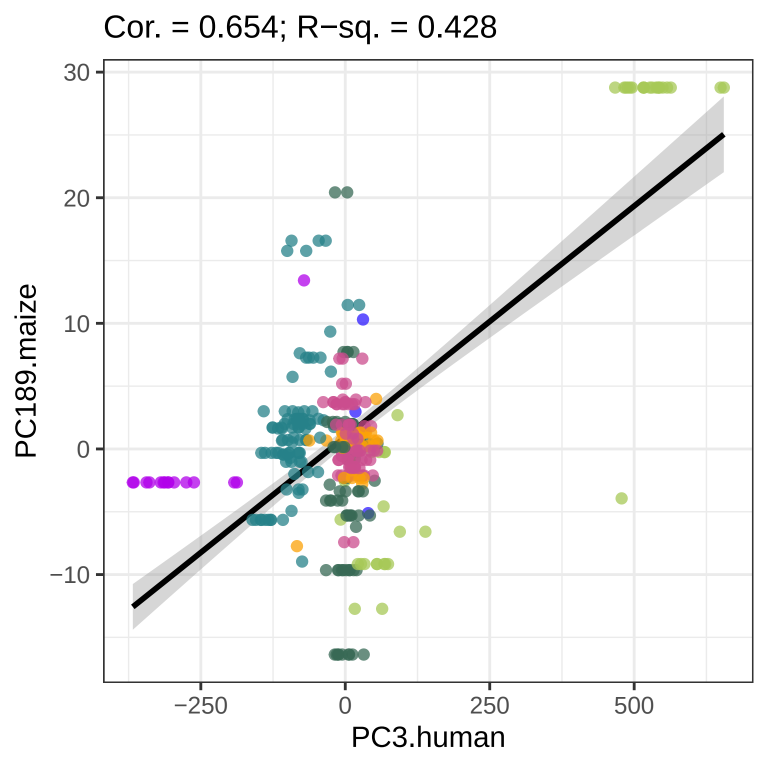

Cor. = 0.654; R−sq. = 0.428
30
20
10
0
−10
PC189.maize
−250
0
250
500
PC3.human

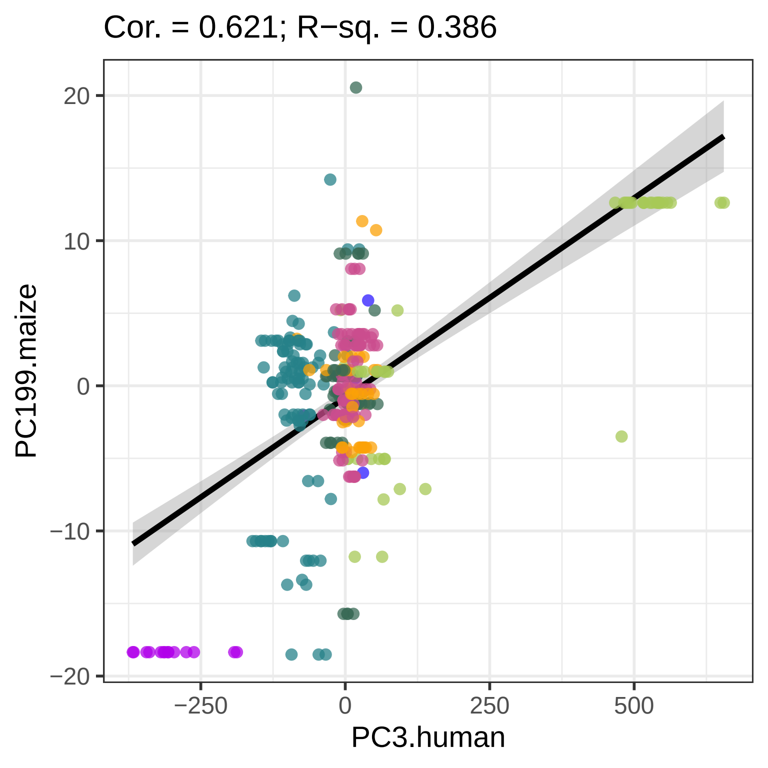

Cor. = 0.621; R−sq. = 0.386
20
10
0
−10
−20
PC199.maize
−250
0
250
500
PC3.human
