## Supplemental Methods for "Maize genetic diversity is largely unstructured by human ethnolinguistic diversity in its center of origin"

### Supplemental text

#### Linguistic polygon range filtering

Previous anthropological and linguistic work were used to identify alternate language names, dialect regions, and resolve other issues for Mayan (Law 2020; Barrett 2008), Uto-Aztecan (Haugen *et al.* 2020), and Otomanguean (Campbell 1997, 2017) datasets. For example, the map polygon "Michoacán Nahuatl" was paired to the village-defined ASJP language "Nahuatl Pomaro Aquila" based on previous linguistic work that described Michoacán Nahuatl as having the majority of its speakers residing in the Aquila and Apatzingán Pomaro municipalities (Campbell 1997).

Common issues included more than one name being used to describe one language, a single language being split into multiple dialects in map ranges or the ASJP dataset, or language dialects defined differently across datasets (including but not limited to: cardinal direction relative to center of language-speaking region, village name, regional name). In the case that multiple potential ASJP languages map to a single language polygon, a grouping is made that encompasses multiple ASJP languages or dialects. For instance, the Otomanguean polygon *Trique* was matched to a larger lexical grouping *OM.MIXTECAN.TRIQUE* consisting of the ASJP languages *TRIQUI_CHICAHUAXTLA*, *TRIQUI_COPALA*, and *TRIQUI_SAN_MARTIN_ITUNYOSO*. For language polygons with specific geographic dialect descriptors (*e.g.* Southern Core Zapotec), ASJP languages with geographic place names (*e.g.* village names) were attributed to these polygons based on the overlap of these place names and the polygon shapefile. This methodology was used most prevalently for large sub-families such as Nahuatl (Uto-Aztecan) and Zapotecan (Otomanguean).

Language polygons that did not have any overlap with existing maize data (see **Maize sampling and genotyping**) were not included. If a language could not be conclusively paired, or separate language names were inconsistently considered to be the same across literature, the polygon and its associated accessions were not included in our study. Some language polygons in both the Haynie and NL range maps were removed based on an inability to match specific ASJP languages to the polygon's range (Northern Alta Mixtec, Northern Baja Mixtec, Subtiaba, Western Zapotec, Jalisco Otomí, Eastern Highland Otomí, and Otomían) and languages that are largely considered extinct and have ranges with little confidence (Zacateco, Cascan, Tecuexe, Uzá).

We validated the polygons of both the Haynie and NL maps by checking whether centroids defined by Glottolog (Forkel and Hammarström 2022) were located within their respective map polygons. In total for Native Land polygons, 11/41, 20/48, and 4/23 languages did not have full overlap with Glottolog centroids for Mayan, Uto-Aztecan, and Otomanguean language families, respectively. For Haynie polygons, 3/21, 7/20, 10/32 language polygons did not have full overlap for Mayan, Uto-Aztecan, and Otomanguean language family Glottolog centroids, respectively. However, of these discrepancies in the Haynie dataset: All Mayan language centroids lay within polygons with the exception of

Chortí and Tz'utujil, which lay near the borders, and Kekchí, which was significantly farther from the centroid. For Uto-Aztecan language polygons, centroids lay near Coatepec Nahuatl, Huichol, Tepecano, and Western Durango Nahuatl, but not near Tahue, Opata, or Eastern Durango Nahuatl. For Otomanguean language polygons, centroids lay near Querétaro Otomí, Atzingo Matlazinca, Eastern Highland Otomí, and Northern Popolocan, but not near Jalisco Otomí, Ixtenco Otomí, Mangue, Michoacán Mazahua, San Francisco Matlatzinca, or Chiapenec. A majority of Otomanguean centroids lay within the Zapotec polygon.

For polygons that were part of the Native Land dataset: For Mayan language polygons, the majority of individual Bats'ik'op polygons did not correspond to Glottlog centroids, but were instead considered as part of the larger Tzotzil subgrouping, which did overlap with a centroid. Additionally, the boundaries of the Mochó polygon did not overlap with its centroid, but was close by. Chontal, Chontal de Guerrero - Tuxteco, and Qyool Mam did not have nearby centroids. For Uto-Aztecan language polygons, most regional Nahuatl dialects that did not overlap were considerably close to centroids, with the exceptions of Huasteca Nahuatl, Coyutecos Nahuatl, Nahuatl de occidente and Nahuatl bajo de occidente being considerably far from potential centroids. In addition, Totorames and Jiak Noki (Yaqui) lay near a centroid on their polygon borders. Ópata (Eudebe and Hobas) polygons additionally did not lay near potential centroids. For Otomanguean language polygons, Mazahua de occidente, Otomí del oeste and Otomí de Ixtenco lay close to, but did not overlap with a centroid. Similar to that of the Haynie polygons, the Chiapaneco centroid did not lay near its corrresponding centroid. Thus, we found that most languages corresponded roughly geographically to an existing Glottolog reference centroid.
